# Mutational impact and signature of ionizing radiation

**DOI:** 10.1101/2021.01.12.426324

**Authors:** Jeonghwan Youk, Hyun Woo Kwon, Joonoh Lim, Eunji Kim, Ryul Kim, Seongyeol Park, Kijong Yi, Sara Jeon, Jinwook Choi, Hyelin Na, Eun-Seok Lee, Young-Won Cho, Dong-Wook Min, Hyojin Kim, Yeong-Rok Kang, Si Ho Choi, Min Ji Bae, Chang Geun Lee, Joon-Goon Kim, Young Seo Kim, Dong Soo Lee, Tae You Kim, Taeyun Ku, Su Yeon Kim, Joo-Hyeon Lee, Bon-Kyoung Koo, Hyunsook Lee, On Vox Yi, Eon Chul Han, Ji Hyun Chang, Kyung Su Kim, Tae Gen Son, Young Seok Ju

**Affiliations:** Graduate School of Medical Science and Engineering, Korea Advanced Institute of Science and Technology (KAIST), Daejeon 34141, Republic of Korea; Department of Molecular Medicine and Biopharmaceutical Sciences, Graduate School of Convergence Science and Technology, Seoul National University, Seoul, Republic of Korea; Department of Nuclear Medicine, Korea University College of Medicine, Seoul, Republic of Korea; Department of Radiation Oncology, Korea Institute of Radiological and Medical Sciences, Seoul, Republic of Korea; Department of Radiation Oncology, Seoul National University College of Medicine, Seoul, Republic of Korea; Department of Biological Sciences & IMBG, Seoul National University. 1 Gwanak-ro, Gwanak-gu, Seoul 08826, Republic of Korea; Wellcome - MRC Cambridge Stem Cell Institute, Jeffrey Cheah Biomedical Centre, University of Cambridge, Cambridge, UK; Department of Physiology, Development and Neuroscience, University of Cambridge, Cambridge, UK; Institute of Molecular Biotechnology of the Austrian Academy of Sciences (IMBA), Vienna Biocenter (VBC), Dr. Bohr-Gasse 3, 1030 Vienna, Austria; Research Center, Dongnam Institute of Radiological and Medical Science, Busan, Republic of Korea; Cancer Research Institute, Seoul National University, Seoul, Republic of Korea; KI for Health Science and Technology, KAIST, Daejeon 34141, Republic of Korea; Department of Nuclear Medicine, Seoul National University College of Medicine, Seoul, Republic of Korea; Department of Internal Medicine, Seoul National University Hospital, Seoul, Republic of Korea; Department of Breast Surgery, Dongnam Institute of Radiological and Medical Science, Busan, Republic of Korea; Department of Surgery, Dongnam Institute of Radiological and Medical Science, Busan, Republic of Korea; Department of Radiation Oncology, Seoul National University Hospital, Seoul, Republic of Korea; Department of Radiation Oncology, Dongnam Institute of Radiological and Medical Science, Busan, Republic of Korea; Department of Radiation Oncology, Ewha Womans University College of Medicine, Seoul, Republic of Korea

## Abstract

Whole-genome sequencing (WGS) of human tumors and normal cells exposed to various carcinogens has revealed distinct mutational patterns that provide deep insights into the DNA damage and repair processes. Although ionizing radiation (IR) is conventionally known as a strong carcinogen, its genome-wide mutational impacts have not been comprehensively investigated at the single-nucleotide level. Here, we explored the mutational landscape of normal single-cells after exposure to the various levels of IR. On average, 1 Gy of IR exposure generated ∼16 mutational events with a spectrum consisting of predominantly small nucleotide deletions and a few characteristic structural variations. In ∼30% of the post-irradiated cells, complex genomic rearrangements, such as *chromoplexy, chromothripsis*, and breakage-fusion-bridge cycles, were resulted, indicating the stochastic and chaotic nature of DNA repair in the presence of the massive number of concurrent DNA double-strand breaks. These mutational signatures were confirmed in the genomes of 22 IR-induced secondary malignancies. With high-resolution genomic snapshots of irradiated cells, our findings provide deep insights into how IR-induced DNA damage and subsequent repair processes operate in mammalian cells.

## Main

Somatic mutations are the most common cause of cancer. These mutations are caused by both exogenous and endogenous mutational processes that are operative in the cell lineages of life since the fertilized egg^1^. Each mutational process is a combination of specific DNA damage and repair mechanisms and generates a definite pattern of mutations, known as ‘mutational signature’^2^. Recent large-scale genome sequencing of multiple cancer types has revealed more than 60 mutational signatures, which provided deep insights into the known and unknown etiologies of the common mutational processes^3^. In addition, experimental exposure of human cells to many agents has enabled examining their direct mutational impact on human cells. Of many different mutational types, base substitutions and small insertions and deletions (indels) have been predominantly investigated because these mutations can be efficiently detected in sequencing. Other classes of mutations, such as copy number changes and genomic rearrangements, have been less widely scrutinized.

Ionizing radiation (IR) is a well-known strong carcinogen that causes irreversible changes in cellular genomes^4^. Despite the frequent exposure to environmental (such as radon gas) and medical radiations, the comprehensive mutational impact and signature of IR-exposure remain elusive Conventional cytogenetic techniques allowed only a rough sketch of chromosomal aberrations resulting from DNA double-strand breaks (DSBs) caused by IR^5,6^. Recent advances in DNA sequencing enabled more sensitive and precise detection of IR-induced mutations in mouse germlines, induced pluripotent stem cells, and radiation-induced tumors^7-10^. However, the insufficient sample size, confounding factors in the experimental systems, and point-mutation-oriented genome analyses limited a comprehensive and quantitative examination of the IR-induced mutagenesis.

For a more comprehensive evaluation of IR’s mutagenic impact, we investigated whole-genomes of irradiated organoids established from various tissue types, having the advantages of being normal, fast-growing, and easy to clone (**Fig. 1a**). Irradiations were conducted by three different approaches that complement each other with various levels of gamma-ray (up to 20 Gy) emitted from Caesium-137/Cobalt-60 or x-ray (up to 60 Gy) from medical linear accelerators (**Fig. 1a; Extended Data Figs. 1a-1c)**. These post-irradiated single-cells were clonally expanded to > ∼100,000 cells using organoid culture technique to perform PCR-free whole-genome sequencing, which markedly minimizes artifacts caused by conventional whole-genome-amplification based methods^11,12^. Overall, we established and explored 156 clonal organoids in 10 different tissues (**Extended Data Fig. 2; Supplementary Tables 1, 2**). Additionally, we further analyzed 22 whole-genomes of IR-induced secondary malignancies (mostly sarcomas), which were extensively irradiated from the medical procedures (**Fig. 1a, Extended Data Fig. 1d**).

**Fig 1.**
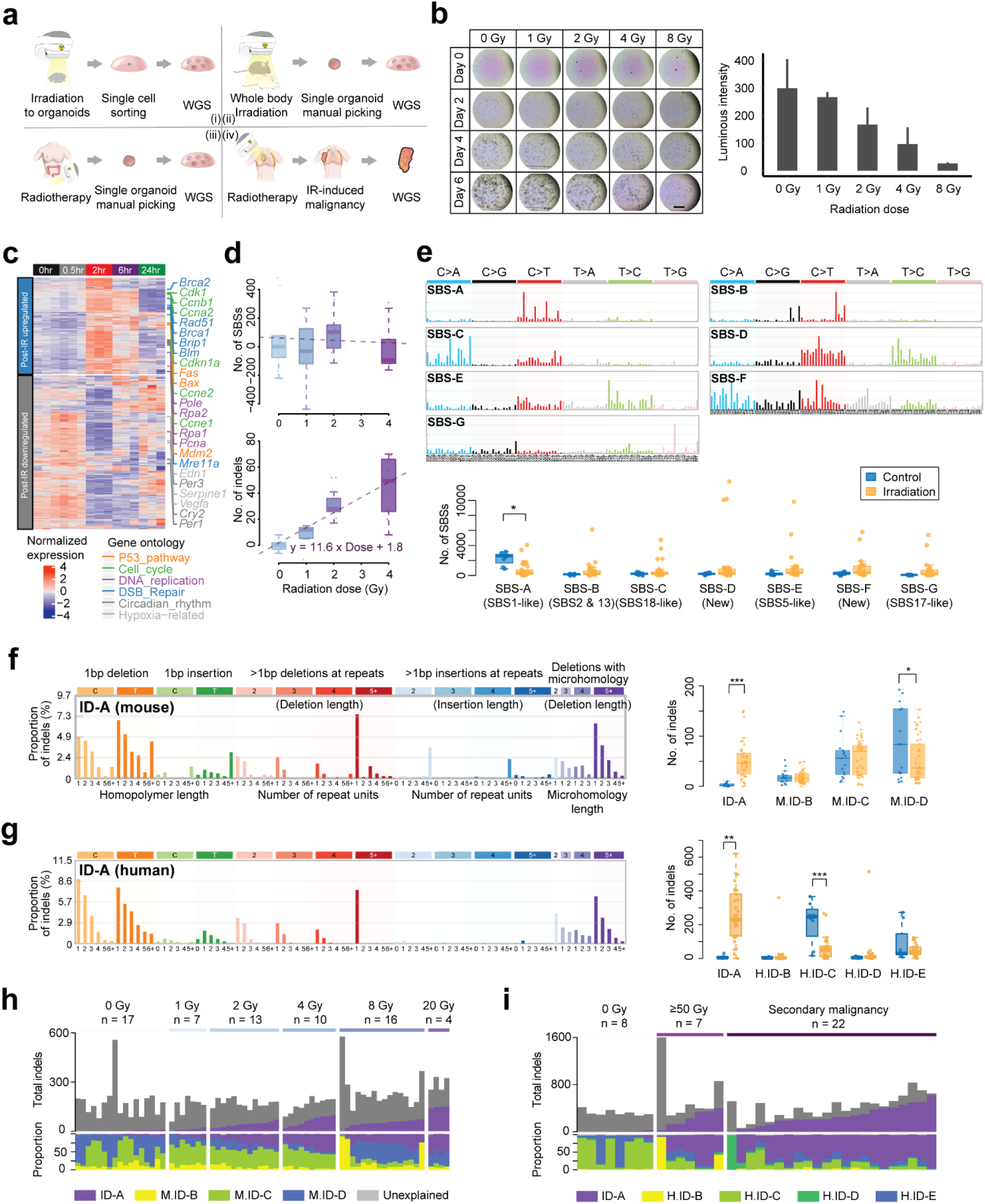
Phenotypic, transcriptomic, and genomic alterations in the irradiated cells. a. Experimental design for detecting IR-induced mutations in single-cells; (i) *in vitro* organoid irradiation (**Extended Data Fig. 1a**), (ii) whole-body irradiation (**Extended Data Fig. 1b**), (iii) human tissue irradiation (**Extended Data Fig. 1c**) followed by clonalization, and (iv) IR-induced secondary malignancies (**Extended Data Fig. 1d**). Whole-genome sequencing was conducted for these clones; WGS: whole genome sequencing; IR: ionizing radiation. b. Organoid-forming efficiencies of post-irradiated cells. Scale bar, 2.5 mm. Error bars represent SEM. n = 3. c. Gene expression changes in cells exposed to 2 Gy irradiation for 24 hrs. Genes in the p53 pathway, cell cycle, DNA replication, and double strand breakage pair pathways are colored orange, green, purple, and blue, respectively. d. Mutational burden of single base substitutions (SBSs; top) and indels (bottom) observed in post-irradiated cells, normalized to that of the non-irradiated (0 Gy). e. SBS signatures extracted from human organoids and IR-induced secondary malignancies (top). The number of SBSs attributable to each signature in control and irradiated groups is shown (bottom). No SBS signature is associated with IR. f. Mutational spectrum of murine indel signature ID-A (left). The other murine indel signatures are shown in **Extended Data Fig. 4a**. Mouse ID-A is associated with IR (right; *: *P* < 0.05; ***: *P* < 0.001). g. Mutational spectrum of human indel signature ID-A (left). The other human indel signatures are shown in **Extended Data Fig. 4b**. Human ID-A is associated with IR (right; **: *P* < 0.01; ***: *P* < 0.001). h. Stacked bar plot showing absolute number of indels (top) and relative proportion of each indel signature (bottom) observed in murine organoids. i. Stacked bar plot showing indel signatures in the human organoids and radiation associated secondary malignancies.

The adverse impact of IR exposure in normal cellular function was observed from the clonalization of single-cells. The organoid-forming efficiency of irradiated single-cells was negatively correlated with the exposure levels (**Fig. 1b**). Roughly, ∼2 Gy of gamma-ray irradiation reduced the proliferation potential by 50% in the pancreas organoids. RNA-sequencing of post-irradiated organoids revealed that many DNA damage response genes, such as *Brca1/2* (in DNA DSB repair pathway), *Mdm2* (in the p53 pathway), and *Cdkn1a* (in the cell cycle pathway), were acutely upregulated and lasted for a few hours after irradiation (**Fig. 1c**). However, these genes were normalized to the baseline in 24 hours, suggesting that the initial cellular response to the irradiation completes within a day (**Fig. 1c**).

From the whole-genome sequencing of the 156 clonal organoids (including 98 irradiated and 58 controls), we catalogued a total of 968,963 somatic single-base substitutions (SBSs) and 87,154 indels. In theory, these mutations include (1) somatic changes that have been acquired during the individual life, (2) IR-induced mutations, and (3) culture-associated mutations that have been accumulated during *in vitro* clonalization. In the SBS mutational type, no substantial correlation was observed between the levels of IR-exposure and the number of SBSs in each clone (**Fig. 1d**). In conventional wisdom, intracellular reactive oxygen species (ROS) generated by IR is known to induce DNA damage^13^, which leads to the C:G>A:T base substitutions defined as SBS signature 18^3^. To assess the ROS-induced mutations more specifically, we decomposed the base substitutions in each cell into mutational signatures^14^ (**Fig. 1e; Supplementary Tables 3**,**4; methods**). However, the mutational burden attributable to SBS-C (equivalent to COSMIC SBS18) was not substantially enriched in the irradiated cells, suggesting that the impact of irradiation is insignificant for the SBS type mutations (**Fig. 1e, Extended Data Fig. 3**). Rather unexpected was that the number of clock-like mutations attributable to SBS-A (equivalent to COSMIC SBS1)^15^ was lower in the irradiated human samples (**Fig. 1e**). It can be explained by tissue donors’ age difference between the irradiated and non-irradiated groups (median age = 75 in non-irradiated and 51 in irradiated samples; *P* = 0.037).

In contrast to SBS, indel type mutations showed a strong association with the levels of IR-exposure (**Fig. 1d**). Of the indel signatures decomposed from the list of somatic indels (**Extended Data Figs. 4a, 4b**; **Supplementary Tables 5, 6**), ID-A was exclusively observed from the irradiated clones (**Figs. 1f, 1g**) and increased in proportion to the IR-exposure levels (**Figs. 1h, 1i**). The indel signature was characterized by (i) 1bp deletion at non- or short-homopolymer sequences and (ii) ≥5 bp deletions. The ≥5 bp deletions showed short overlapping microhomology in the breakpoint junctions of more frequently than random expectation (**Extended Data Fig. 4c**). This signature is similar to COSMIC ID8, which had been suggested for mutations caused by errors in non-homologous end joining (NHEJ) of the double-strand breaks of DNA^3^ (**Extended Data Fig. 4d)**.

As expected, the indel signature was clearly observed from the whole-genomes of radiation-induced secondary malignancies (**Fig. 1i**). In these samples, >50% of the total indels discovered were attributable to the indel mutational signature, ID-A (**Supplementary Tables 7, 8)**. Compared with whole-genomes of 71 primary sarcomas as control^16^, the burden of ID-A indels was much higher in the IR-induced secondary malignancies: ID-A accounts for more than 1/3 of indels in 20 cases of the secondary malignancies in contrast to 6 cases in 71 primary sarcomas (92% vs. 8%; *P* < 0.001; **Supplementary Tables 8**).

Ionizing radiation also caused characteristic structural variations (**Fig. 2a**). The balanced-inversion type was significantly associated with IR, as reported in the literature^10^. The balanced-inversion type was observed in more than a half of the irradiated samples, with a higher burden in the clones of higher exposure levels, whereas none were found in the non-irradiated organoids (64%, 46%, and 0% of the clones in ≥2 Gy, 1 Gy, and 0 Gy irradiated groups, respectively; *P* < 0.001). In addition, the balanced-translocation type and long-deletion type (≥1 Mb) mutations were almost exclusively observed in the irradiated organoids (balanced translocation: 45%, 8%, 0% of the clones in ≥2 Gy, 1 Gy, and 0 Gy irradiated groups, respectively, *P* < 0.001; long-deletion: 33%, 8%, 2% of the clones in ≥2 Gy, 1 Gy, and 0 Gy irradiated groups, respectively, *P* < 0.001). Extrachromosomal DNA elements, e.g., double minutes^17^, were found (4%) in the highly irradiated samples (≥2 Gy). Some SV types, such as medium-sized deletion (100 bp to 1 Mb), tandem duplication, and templated insertion, were observed in the non-irradiated clones with substantial frequency, suggesting that these are non-specific to irradiation and can be acquired during normal aging or during organoid culture.

**Fig 2.**
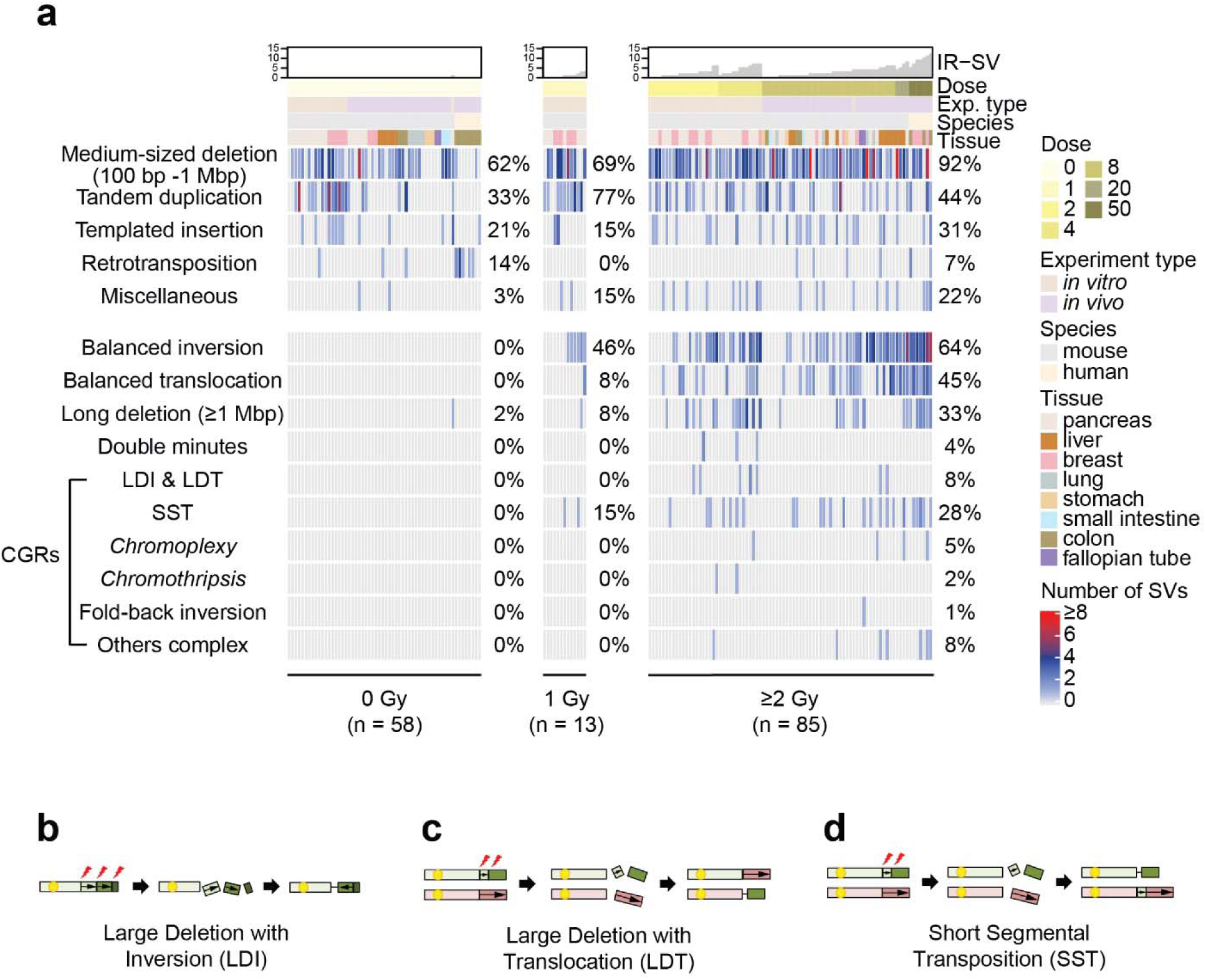
Signatures of IR-induced structural variation. a. Heatmap showing IR-induced structural variations (SVs). Each row indicates SV type, with each column representing the number of SVs observed in each sample. Samples are classified into three groups (0, 1, and ≥2 Gy (n = 58, 13, and 85, respectively)). SV types frequently seen in the non-irradiated group are represented in top panel. SV types specifically observed in the post-irradiated organoids are represented in bottom rows; LDI: large deletion with inversion; LDT: large deletion with translocation; SST: short segmental transposition; IR-SV: IR-induced structural variation. b. Schematic diagram for LDI. c. Schematic diagram for LDT. d. Schematic diagram for SST.

From our clones, we identified 64 SV events consisting of ≥3 interconnected double-strand DNA breaks defined as complex genomic rearrangements (CGRs) (**Fig. 2a**). The simplest patterns of the CGRs were the large deletions combined with local inversions or translocations (referred to as LDIs and LDTs, respectively; **Figs. 2b, c**). LDI- and LDT-type mutations were exclusively found in the irradiated organoids with a frequency of ∼5% and ∼2%, respectively. In addition, we found another type of CGRs, characterized by an insertion of a short DNA fragment scissored-out from a genomic location (hereafter referred to as short segmental transpositions (SSTs)) as seen in the cut-and-paste mobilization of DNA transposons (**Figs. 2d, 3a; Extended Data Fig. 5**). The length of the DNA fragment typically ranged from 100 bp to 1000 bp. All 29 SST events discovered in this study were exclusively observed in the irradiated group (27% vs. 0% of the clones in irradiated and control groups, respectively; *P* < 0.001; **Fig. 2a**). Intriguingly, 11 SST events were found in the radiation-induced secondary malignancies, with a 10-fold higher frequency than in general primary cancers^16,18^ (*P* < 0.001; **Fig. 3b**).

**Fig 3.**
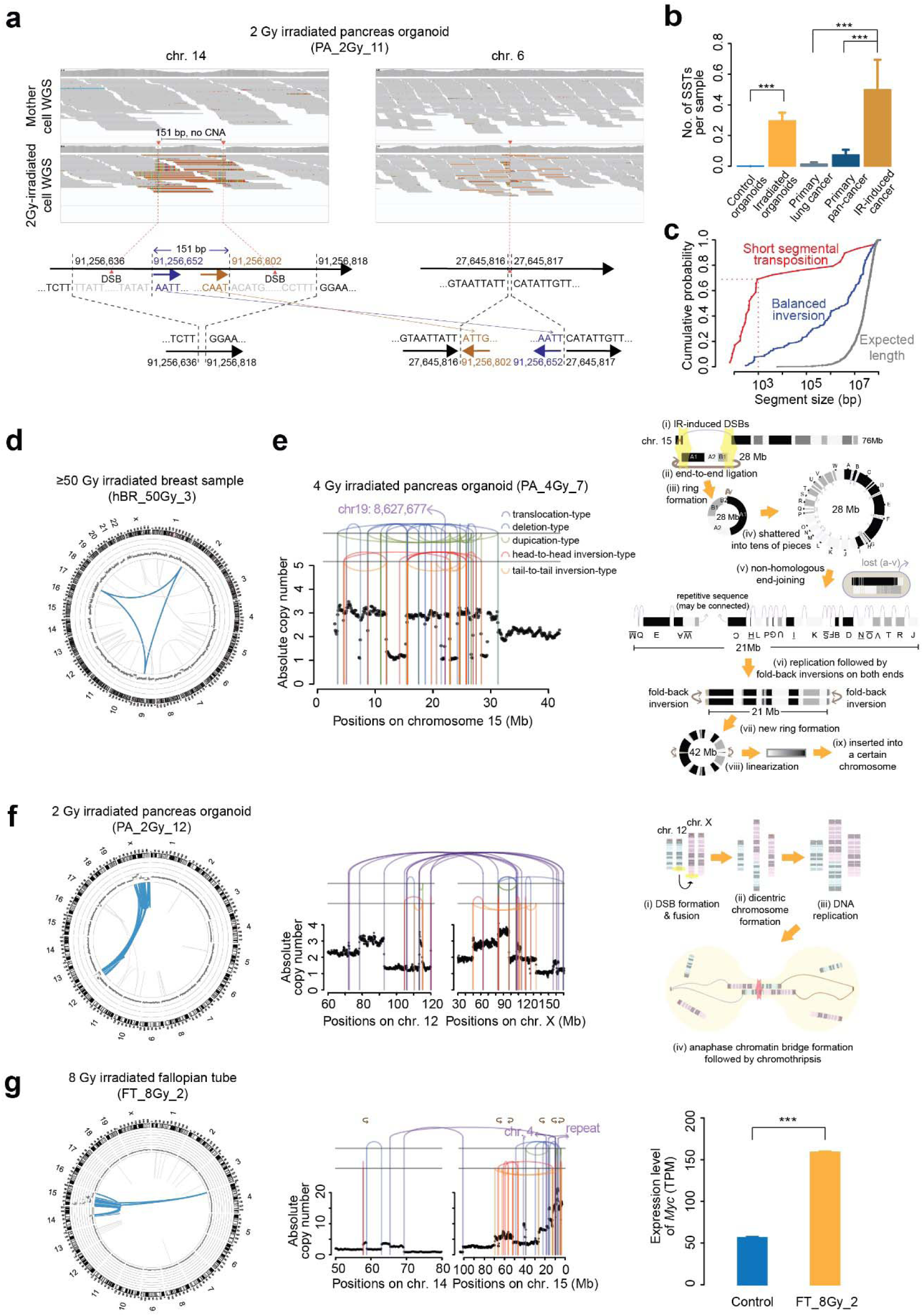
Complex genomic rearrangements induced by ionizing radiation. a. An example of the short segmental transposition (SST) event. A 151 bp segment in chromosome 14 was cut in the original place and then inserted into the middle of chromosome 6. Notably, it is a copy number neutral event; WGS: whole genome sequencing; CNA: copy number alteration; DSB: double strand break. b. Frequency of SST events in non-irradiated organoids (n = 58), irradiated organoids (n = 98), primary lung cancer^18^ (n = 138), primary pan-cancer^16^ (n = 231), and IR-induced secondary cancer (n = 22). ***: *P* < 0.001. Error bars represent SEM. c. Segmental length distribution of SST, balanced inversion, and in-silico expectation of two breakpoints in a 2 Gb chromosome (***: *P* < 0.001). d. A *chromoplexy* event (blue lines) found in a post-irradiated human organoid (hBR_50Gy_3). From outer to inner layer, the circos plot depicts chromosome ideogram, scatter plots for copy number status (one dot represents mean copy number status of 100 Kb; each space between two lines represents 2 copy number differences), and lines for structural variations. e. *Chromothripsis* found in a post-irradiated human organoid (PA_4Gy_7). Patterns of rearrangements and copy number status in chromosome 15p (left). One simple scenario that can generate the event (right). f. *Chromothripsis* (blue lines) localized in chromosomes 12 and X found in a post-irradiated organoid (PA_2Gy_12; left). Patterns of rearrangements and copy number status (middle). One simple scenario that can generate the event (right). g. Fold-back inversions through breakage-fusion-bridge cycles (BFB cycles; blue lines) found in a post-irradiated human organoid (FT_8Gy_2; left). Patterns of rearrangements and copy numbers (middle). The difference of *Myc* expression in control and irradiated organoids (FT_8Gy_2). Error bars represent SEM. n = 3; TPM: transcripts per million.

The balanced-inversion type rearrangement can be considered a special type of SST, in which a broken DNA fragment is inserted in its original site with inverted orientation. Balanced-inversions and SSTs showed significant differences in the length of broken DNA fragments (5.2 Mb vs. 422 bp in median, respectively; *P* < 0.001; **Fig. 3c**). We speculate that shorter DNA fragments are more mobile and can migrate to a distant position in the nucleus before rejoining into chromosomal structures by DNA repair mechanisms.

Of the 85 organoids irradiated with ≥2 Gy, seven (8.2%) harbored highly catastrophic CGRs, including *chromoplexy, chromothripsis*, and SVs by breakage-fusion-bridge (BFB) cycles^19^. Four of them had *chromoplexy* events, also known as “closed-chain” multi-chromosomal balanced translocations. *Chromoplexy* frequently shapes oncogene rearrangements in prostate and lung cancers^18,20^ (**Fig. 3d, Extended Data Figs. 6a-c**). Despite its functional importance, the mutagenic origin of *chromoplexy* has been unknown. We speculate that *chromoplexy* can be structured in an irradiated cell when IR-induced, concurrent DNA breaks of multiple chromosomes are stochastically ligated with an erroneous configuration. Our findings indicate that IR exposure can be an important origin for *chromoplexy*, although its absolute epidemiological contribution is uncertain.

*Chromothripsis* events, characterized by extensive chromosome shattering and reassembly^21^, were also observed in two post-irradiated organoids. For instance, a clone exposed to 4 Gy radiation (PA_4Gy_7) harbored localized massive genomic rearrangements in chromosome 15, consisting of ≥47 breakpoints and an oscillation of DNA copy number between one and three (**Fig. 3e, Extended Data Fig. 6d**). Through careful reconstruction of the complex event, we found that the vast majority of the breaks were generated by errors in the IR-induced DNA repair processes rather than by direct physical damages. In the simplest scenario that might construct the *chromothripsis* event, IR generated only two breakpoints in the chromosome, leading to the excision of 28Mb-long DNA fragment followed by the formation of an intermediate ring structure by end-to-end ligation of the fragment from the erroneous DNA repair (**Fig. 3e**). Due to the absence of a centromere and telomeres, the ring structure is structurally unstable and is vulnerable to be shattered and reassembled in the subsequent cell divisions. These events result in extensive rearrangements^22^ until the structure is either eradicated or stabilized by re-insertion into the chromosome. In another clone with 2 Gy exposure (PA_2Gy_12), two chromosomes (chromosomes 12 and X) were jointly involved in a *chromothripsis* event (**Fig. 3f**). C*hromothripsis* can involve two chromosomes if two chromosomes are fused end-to-end with having two centromeres (dicentric chromosome). The subsequent cell division pulls the two centromeres to opposite poles and results in an anaphase chromatin bridge and massive shattering^23,24^. Therefore, we speculated that IR-exposure caused two chromosomes’ telomeric losses, followed by a misplaced end-to-end DNA repair. Supporting the mechanism, we observed a localized hypermutation, known as *kataegis*, in the vicinity of the breakpoints (**Extended Data Fig. 6e**), which is an imprint of the chromatin bridge^23^. We also observed a post-irradiated clone (FT_8Gy_2) harboring a structural variation called fold-back inversion ^25^, which is shaped by the breakage-fusion-bridge (BFB) cycle^26^, another type of complex rearrangement typically resulting from an erroneous repair of the dicentric chromosome structure (**Fig. 3g, Extended Data Fig. 6f**). The event amplified *Myc* oncogene to total 6 copies, leading to the transcriptional upregulation of the gene (from 56 to 159 TPMs, *P* < 0.001; **Fig. 3g**). Collectively, our findings suggest that IR-exposure can induce cellular catastrophes by generating unstable intermediate chromosomal structures.

Among the aforementioned SV types, a few types were exclusively observed in the irradiated clones. These include balanced inversion, balanced translocation, SST, *chromoplexy, chromothripsis*, and fold-back inversion events (**Fig. 2a**). To explore the quantitative mutagenic impact of IR, we regarded the SV types as IR-induced SVs (IR-SVs). Irradiation of 2 Gy or higher produced at least one IR-SV event in ∼90% of the cells (76 out of 85). These IR-SVs were distributed in all types of organoids, and any tissue-type specificity was not observed. The number of IR-SVs (**Fig. 4a**) showed a strong linear correlation with the total levels of IR-exposure. Considering the numbers of IR-induced indels (**Fig. 1d**), mutation-free DSB repairs (**methods**), and IR-SVs, we estimated that 1 Gy of gamma irradiation causes 16.1 DSBs in a murine cell (which is equivalent to 18.3 DSBs in human cells) (**Fig. 4b**). The number agrees well with our counting from 3D super-resolution imaging of the post-irradiated cells (**Fig. 4c**) and falls in the range of the rough evaluation in the previous reports^27,28^. When a human cell was exposed by 1 Gy of high-rate IR (i.e., short-term full exposure with ∼0.5 Gy/min), these 18.3 DSBs were comprised of 14.9 for indel mutations, 2.5 for SVs, and ∼1 for seamless ligation. The frequency of the mutation-free DSB ligations was estimated to be as low as 5%. Exposure of cells to the same dose but at a very low rate (∼0.08Gy/day) produced a similar number of overall DSBs, but the density of SVs decreased by 50% (**Extended Data Fig. 7a**), suggesting that the higher number of concurrent DSBs produced by high-rate IR increases the likelihood of the misplaced end-joining between discordant DNA ends.

**Fig 4.**
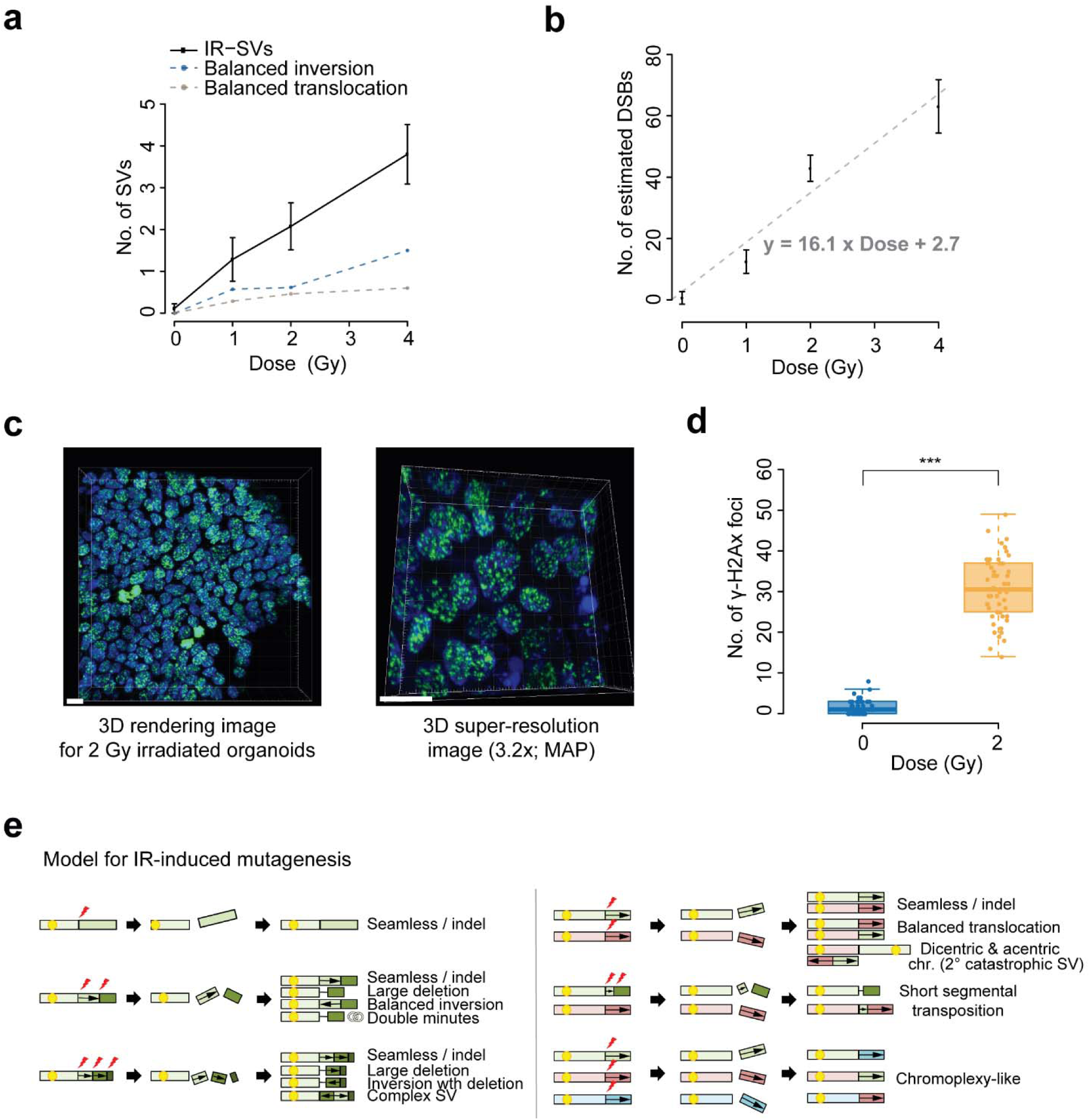
Dose-response curves and patterns of ionizing-radiation-induced variations. a. Number of IR-induced structural variations (IR-SVs) is linearly proportional to IR dose. b. Number of estimated DSBs according to IR dose in irradiated murine pancreas organoids. 1 Gy of IR induces about 16.1 DSBs, Error bars represent SEM. DSB: double-strand breakage. c. Representative image of γ-H2Ax immunofluorescent staining (green) in 2 Gy irradiated murine pancreas organoids (left; 63x). 3.2x expanded γ-H2Ax immunofluorescent staining image (right; 63x). DAPI: blue. MAP: magnified analysis of the proteome. Scale bars, 10 μm. d. Number of γ-H2Ax foci in control and 2 Gy irradiated organoids (1.5 and 30.7 DSBs on average, respectively). Fifty cells in each group were randomly selected and counted (***: *P* < 0.001). e. Schematic illustration of IR-induced mutagenesis model. Consequences of IR-induced breakpoints on one chromosome (left). Consequences of IR-induced breakpoints on multiple chromosomes (right). Unstable chromosome intermediates such as dicentric chromosomes and acentric chromosomes stochastically resulted in catastrophic genomic rearrangements.

Sequence characteristics in the breakpoint junctions provide insights into the mechanism of IR-induced-DSB repair. Indeed, the breakpoint junctions of IR-SVs in irradiated organoids (n = 590) were clearly distinguished from those of non-IR-SVs in non-irradiated organoids (n = 223). For example, the length of breakpoint microhomology was significantly shorter in IR-SVs (1 bp vs. 2 bp in median; *P* < 0.001; **Extended Data Fig. 7b**). In addition, the majority of IR-SV breakpoints (65%) were accompanied by short nucleotide deletions (1-20 bp; **Extended Data Fig. 7c**). Compared with those of non-IR-SVs, the breakpoints of IR-SVs were not significantly associated with local genomic features, such as chromatin status, replication timing, and GC ratio (**Extended Data Fig. 7d**). These collectively suggest that IR induces DSBs more-or-less randomly in the genome and that the non-homologous end joining (NHEJ) mechanism is dominantly responsible for the repair^29^.

To reveal the comprehensive signature of IR-induced mutations, we combined organoid and whole-genome-sequencing technologies. By doing so, we could systematically observe the genome-wide mutational impact of IR, including both the small nucleotide-level (SBS and indels) and the large structural variations on the largest scale ever to the best of our knowledge. We estimated that 1 Gy of IR-exposure generates ∼16 mutations genome-wide. However, the mutational rate should be interpreted more carefully: our experimental system cannot investigate non-dividing cells harboring the massive IR-induced genomic damages or critical defects.

We detected signatures of IR-induced indels and IR-SVs that were observed exclusively in the post-irradiated normal cells. Although our efforts provide an average reference signature by irradiation, genomic locations of DNA double-strand breaks are random and thus genomic architectures after recovery from damage can be highly variable across cells. Depending on the formation timing, spatial proximity, and topological configuration of the IR-induced DSBs, the likelihood for their correct ligation would be highly altered (**Fig. 4e**). When IR-induced DSBs are followed by the formation of unstable chromosomal structures (such as the dicentric chromosome), massive fragmentation of chromosomes can be caused, initiating extensive genomic rearrangements. We observed that ∼10% of clones with ≥2 Gy IR-exposure experienced these catastrophic events.

Our findings directly suggest that the vast majority of apparently normal cells in the field of therapeutic irradiation harbor IR-induced mutations. In typical radiotherapy, millions of normal cells in the adjacent normal tissues are typically exposed to 50 Gy or higher irradiation. Given the mutation rate estimated in the study, hundreds of mutations will be on average acquired per single-cell. Because IR-induced mutation occurs at random genomic positions, some cells will gain functionally critical mutations by chance, such as in cancer-related genes. These mutant cells may reside and proliferate in the tissue, silently surviving for a long time.

In our experimental setting, we irradiated our cells with a few defined conditions, such as by the ^137^Caesium irradiators with a few fixed irradiation rates for conceptual and practical simplicity. However, we do not exclude the possibility of the distinct mutational profiles when cells are exposed to different types of irradiation or under the ultra-high (or low) radiation rate. Therefore, similar analyses of more irradiated clones will likely yield a more comprehensive mutational signature of IR.

## Data availability

Whole-genome and transcriptome sequencing raw data will be deposited on the Sequence Read Archive (SRA) for mouse data and the European Genome Archive (EGA) for human data. Accession IDs are not assigned yet. Organoids established in this study will be available under a material transfer agreement. All data will be provided for review purposes upon reviewers’ request

## Code availability

All in-house scripts used in this study will be posted on Github. All computational codes will be provided for review purposes upon reviewers’ request.

## Acknowledgements

The authors thank Jinwoo Seong and Young-Yun Kong (Seoul national university) for mouse mammary gland acquisition, and Myungsuk Choi (KAIST) and Jong-Yeon Shin (Macrogen Inc.) for technical help. This work was supported by the Suh Kyungbae Foundation (SUHF-18010082 to Y.S.J.); Young Investigator Grants from the Human Frontier Science Program (RGY0071/2018; to Y.S.J. and B.-K.K); the National Research Foundation of Korea (Leading Researcher Program NRF-2020R1A3B2078973 to Y.S.J; for Brain Pool Program NRF-2019H1D3A2A02061168 to Y.S.J. and S.Y.K.; Basic Science Research Program NRF-2020R1I1A1A01072873 to K.S.K., NRF-2017R1A2B4003535 to J.H.C.); the National R & D Program funded by the Ministry of Science, ICT & Future Planning through the Dongnam Institute of Radiological & Medical Sciences (DIRAMS) under grants (50491-2018 and 50491-2019 to T.G.S.); Research grant from Korea University Anam Hospital, Seoul, Republic of Korea (Grant No. O1800741, O2000681 to H.W.K).

## Author contributions

Y.S.J., T.G.S., K.S.K., J.Y., and H.W.K. designed the experiments. H.W.K., T.G.S., H.K., Y.-R.K., C.G.L., and M.J.B. conducted in vitro and in vivo mouse irradiation experiments. K.S.K., O.V.Y., and E.C.H. provided adjacent human normal samples. E.K. and J.H.C. provided secondary malignancy samples. K.S.K., E.K., and J.H.C. oversaw all clinical data collection and curation. J.Y., H.W.K., E.-S.L., and T.G.S. performed organoid culture, clonal expansion, and DNA/RNA extraction, with S.H.C., and D.S.L. providing advice. S.K. (pancreas), J.C. (lung), H.N. (Wnt), J.L. (lung), B.-K. K. (Wnt, colon), and H.L. (pancreas) providing training in organoid culture technologies. H.W.K. performed cell viability analysis and organoid staining with Y.-W. C., D.-W. M., and T.Y.K.. H.W.K., J.-G. K., Y.S.K., and T.K. performed organoid expansion and obtained high-resolution confocal images. J.Y. performed most bioinformatic analysis including alignment, mutation calling, and data curation, with S.P., K.Y., R.K., and Y.S.J. providing advice. J.L. carried out mutational signature analysis. S.Y.K. oversaw statistical analysis. J.Y., H.W.K., J.L., K.S.K., T.G.S., and Y.S.J. wrote the manuscript. All authors finalized the manuscript. Y.S.J. supervised the project.

---

**Extended Data Fig. 1.**
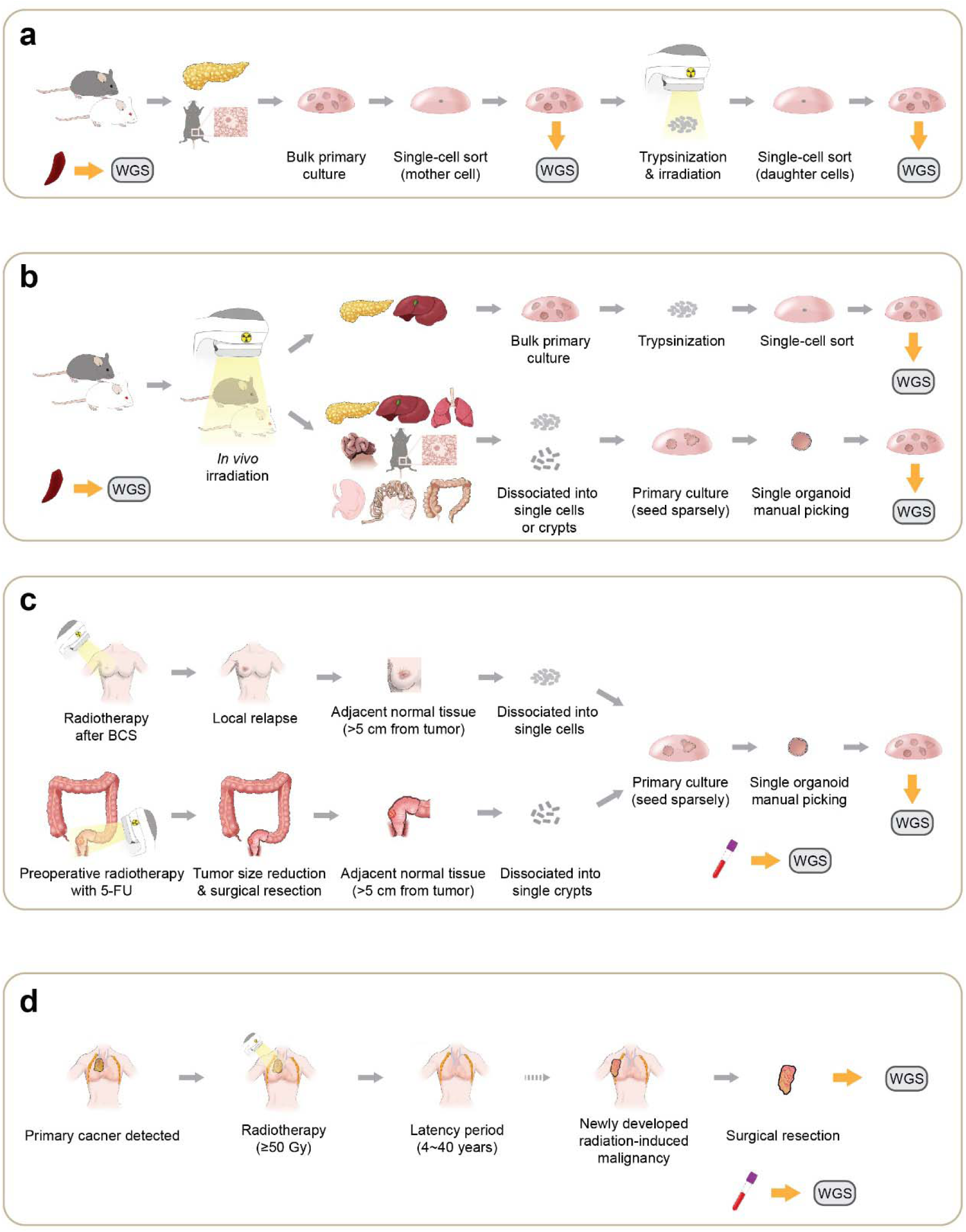
Schematic illustrations of experimental procedures. a. Schematic procedures for *in vitro* organoid irradiation applied to pancreas and breast organoids. Single cells were sorted by fluorescence-activated cell sorter (FACS). Whole-genome sequences of irradiated clones were compared with genome sequences of pre-irradiated mother cells or spleen to isolate somatic mutations. b. Schematic procedures for *in vivo* whole-body irradiation. After irradiation, clonal expansion and whole-genome sequencing followed for detecting IR-mediated somatic mutations. c. Schematic illustration for the collection of human irradiated single cells followed by clonalization and whole-genome sequencing. Whole-genome sequences of matched blood DNA were used as a germline reference; BCS: breast conserving surgery; 5-FU: 5-fluoro uracil. d. Sequencing of radiation-induced secondary malignancies.

**Extended Data Fig. 2.**
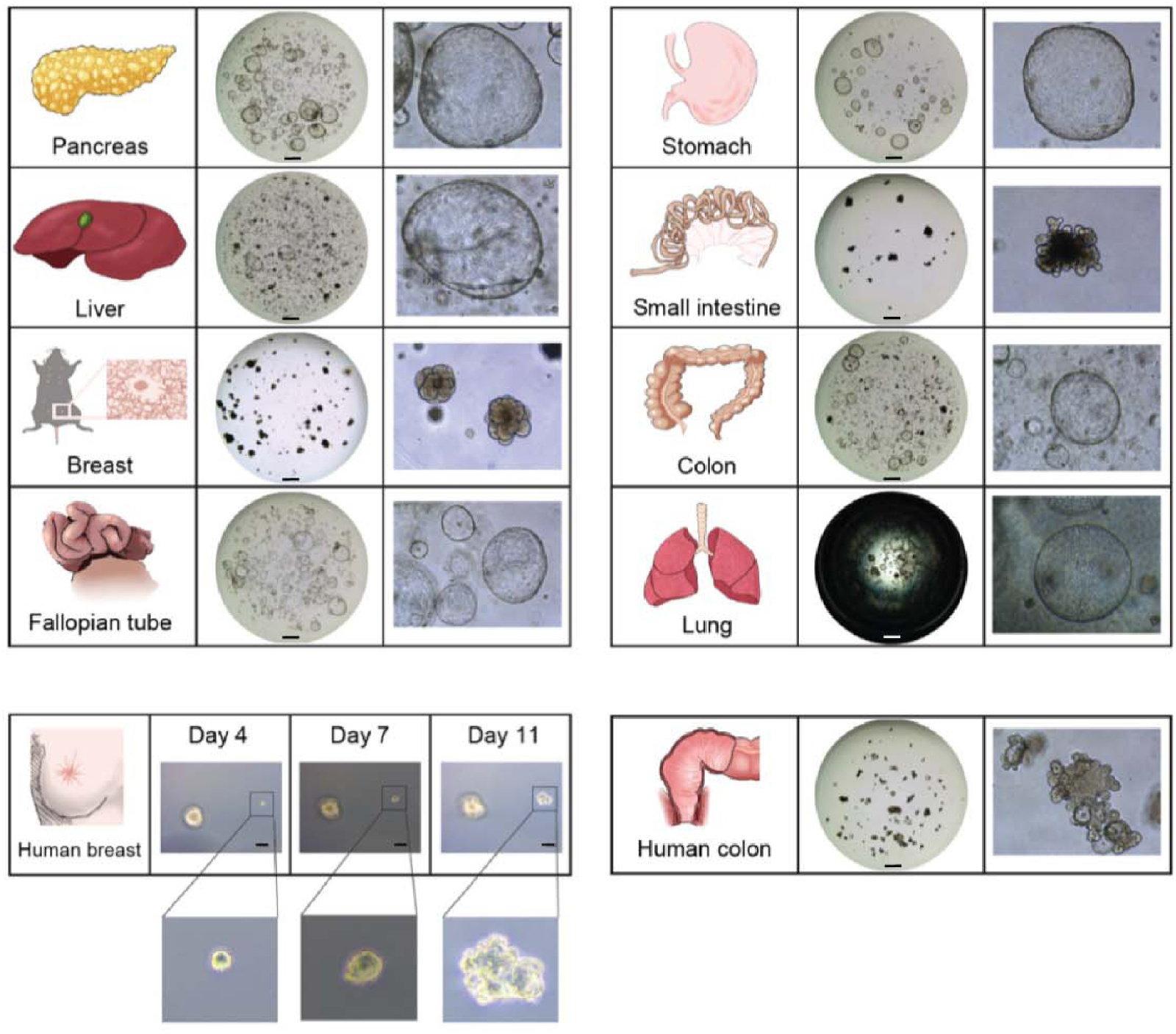
Representative images of organoid types established in this study. Post-irradiated organoids were established from eight murine and two human tissues in the study. Scale bar is 1 mm, except human breast images (20 μm).

**Extended Data Fig. 3.**
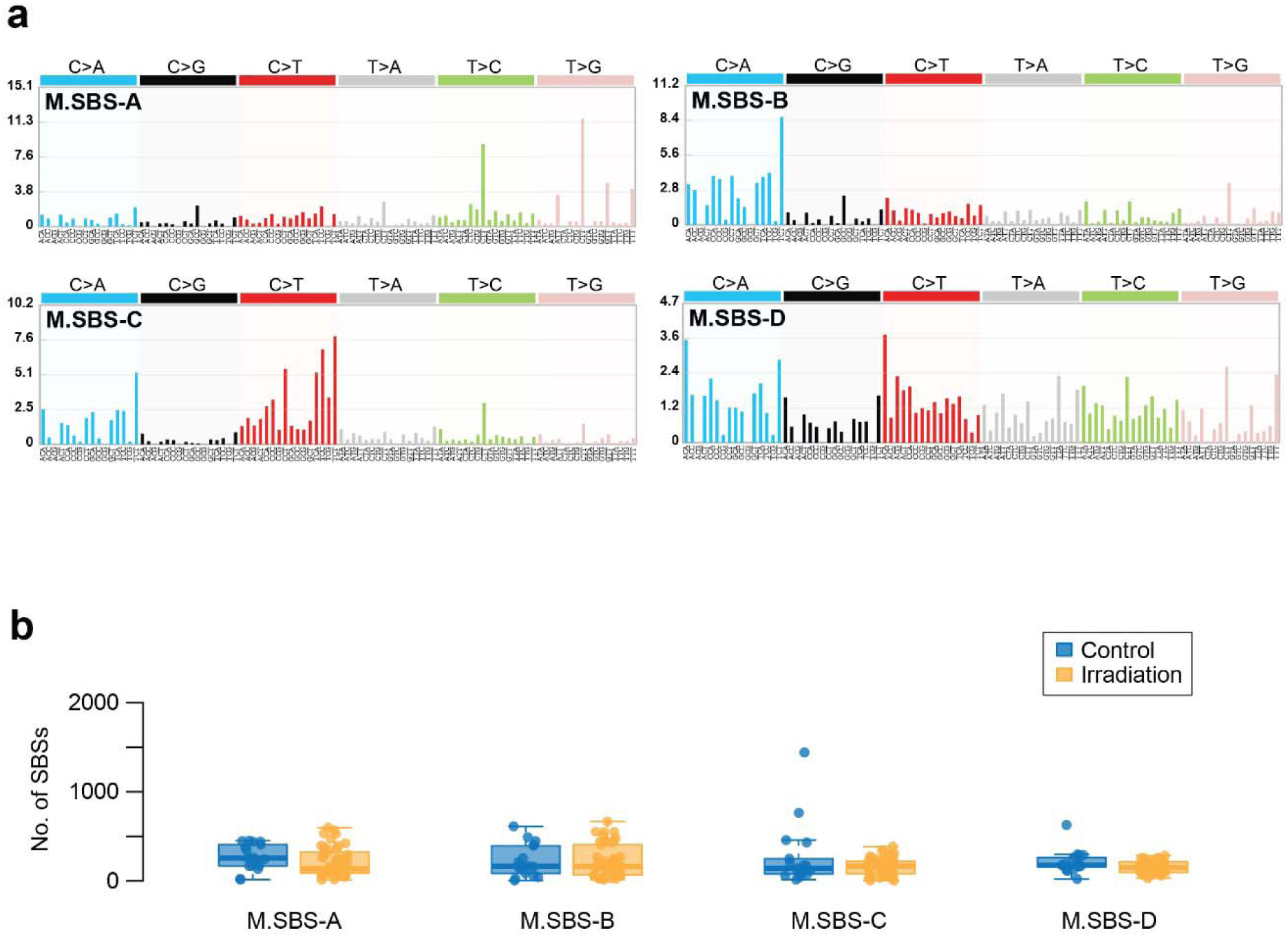
Mutational signatures for somatic single base substitutions (SBSs) identified in murine organoids. a. Four SBS signatures extracted from 67 murine monoclonal organoids. b. Mutational burden attributed to each SBS signature according to irradiation.

**Extended Data Fig. 4.**
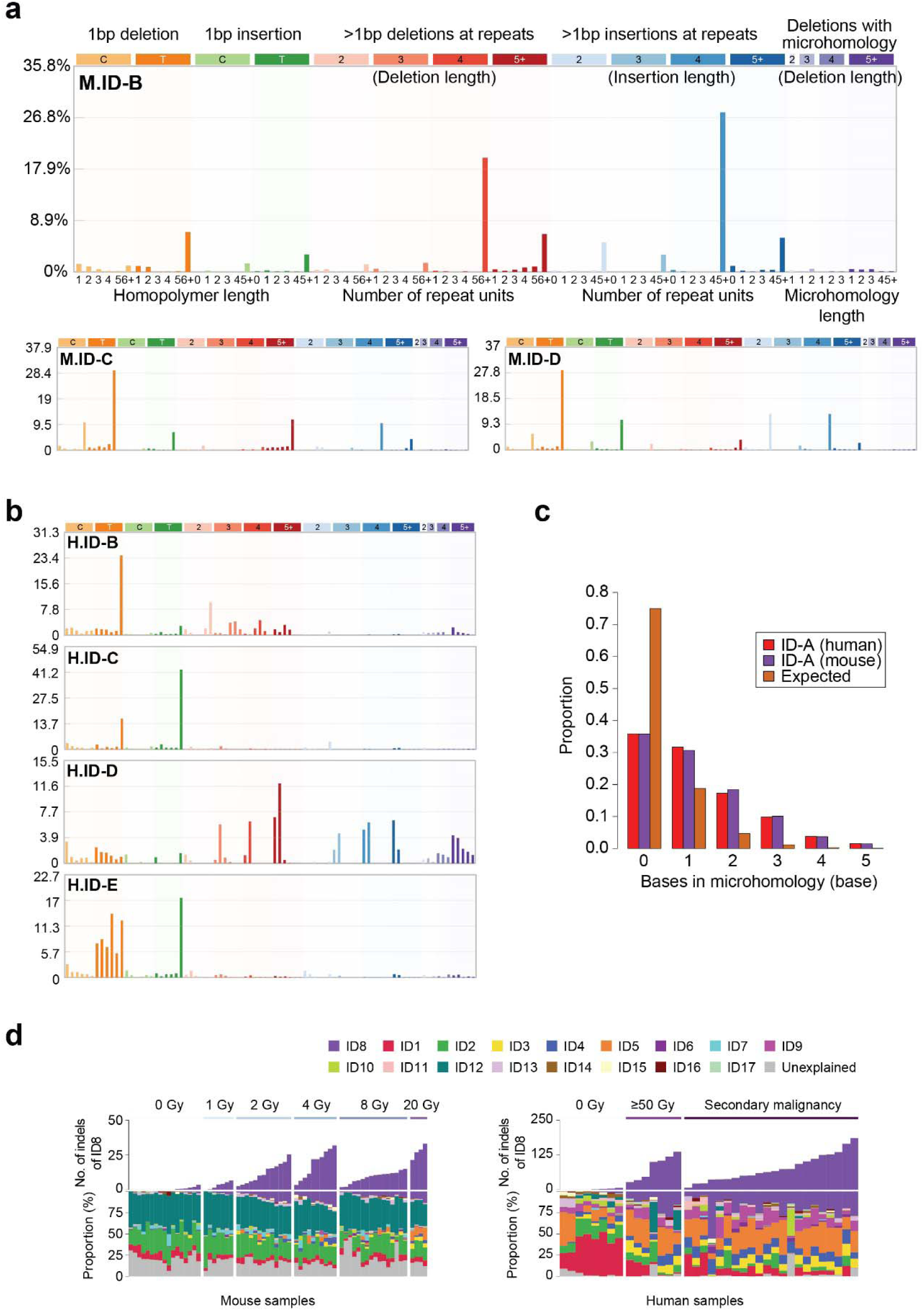
Indel signatures extracted from irradiated samples. a. Mouse indel signatures except mouse ID-A. b. Human indel signatures except human ID-A. c. Sequence microhomologies observed in IR-associated small deletions ≥5 bp. These indels showed higher levels of microhomology than expected by chance. d. COSMIC ID signatures were fitted to the mutational spectra of the control and the irradiated samples. COSMIC ID8 signature (purple) is specifically observed in irradiated samples.

**Extended Data Fig. 5.**
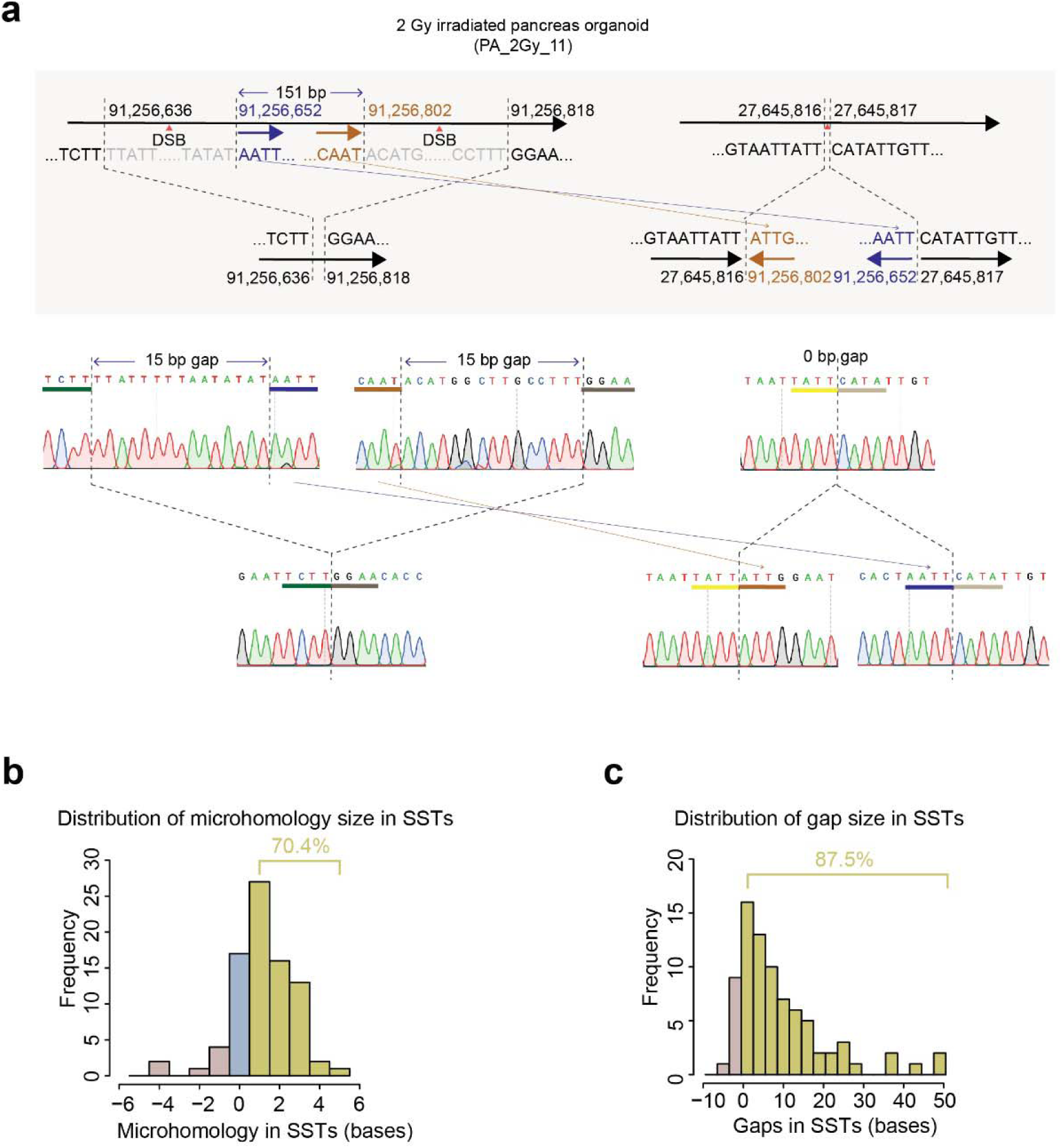
Sequence characteristics of short segmental transpositions (SSTs). a. Sanger validation of an SST case. b. Distribution of microhomology at SST breakpoints. More than 70% of breakpoints show >0 bp microhomology. c. Distribution of gap sizes in breakpoints of SSTs. Most breakpoints (>80%) have short deletions suggestive of end-repairs of DSBs.

**Extended Data Fig. 6.**
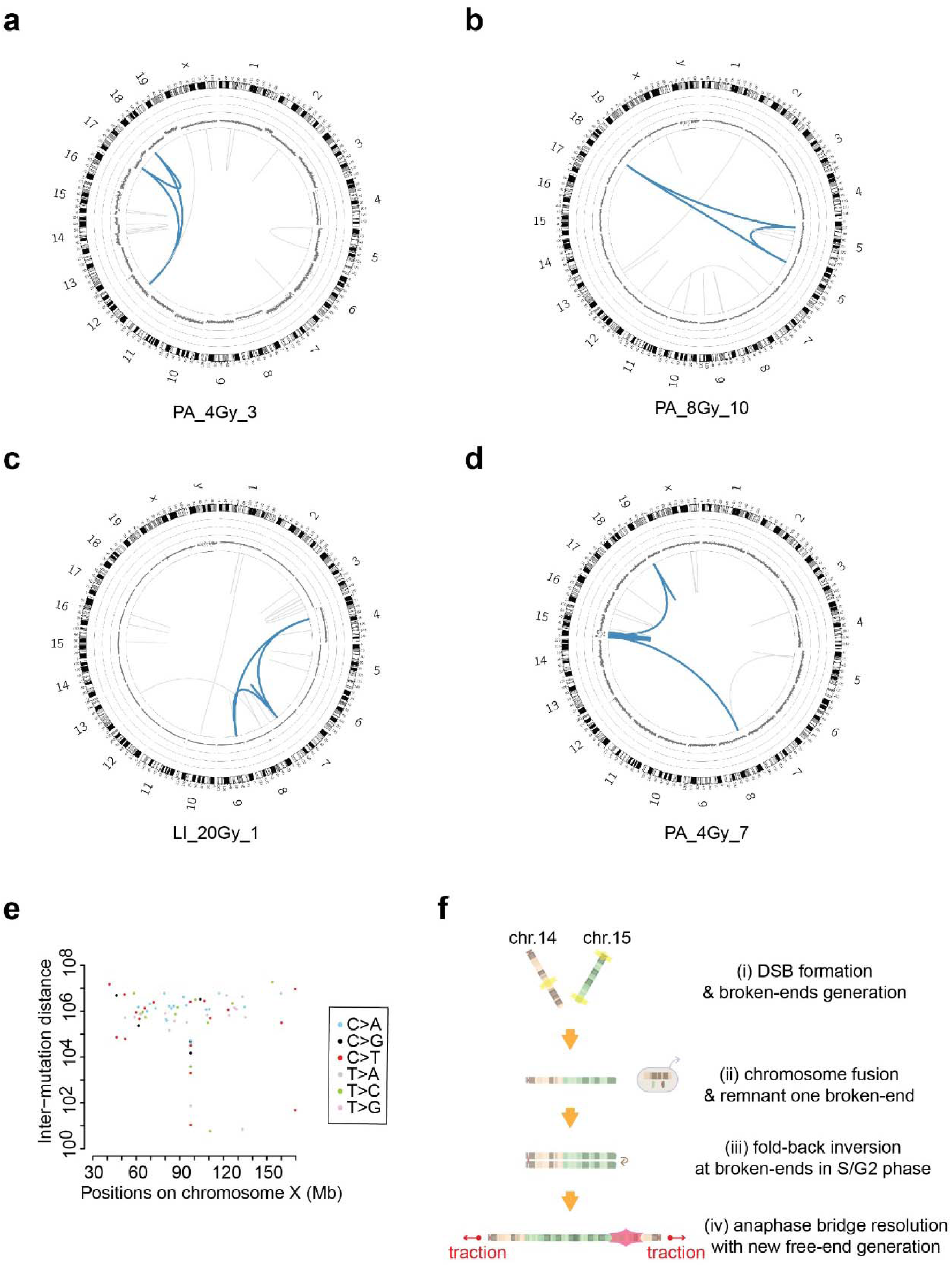
*Chromoplexy* and *chromothripsis* identified in irradiated organoids. a. Circos plot showing *chromoplexy* (blue lines) in a 4 Gy irradiated organoid. b. Circos plot showing *chromoplexy* (blue lines) in an 8 Gy irradiated organoid. c. Circos plot showing *chromoplexy* (blue lines) in a 20 Gy irradiated organoid. d. Circos plot showing *chromothiripsis* (blue lines) on chromosome 16 in a 4 Gy irradiated organoid. The copy number state in the catastrophic segment oscillates between three and one. e. Rainfall plot map demonstrating *kataegis* at chrX:97.5M observed with a *chromothripsis* event in PA_2Gy_12. f. Schematics of a possible scenario for the IR-induced breakage-fusion-bridge cycles occurred in FA_8Gy_2.

**Extended Data Fig. 7.**
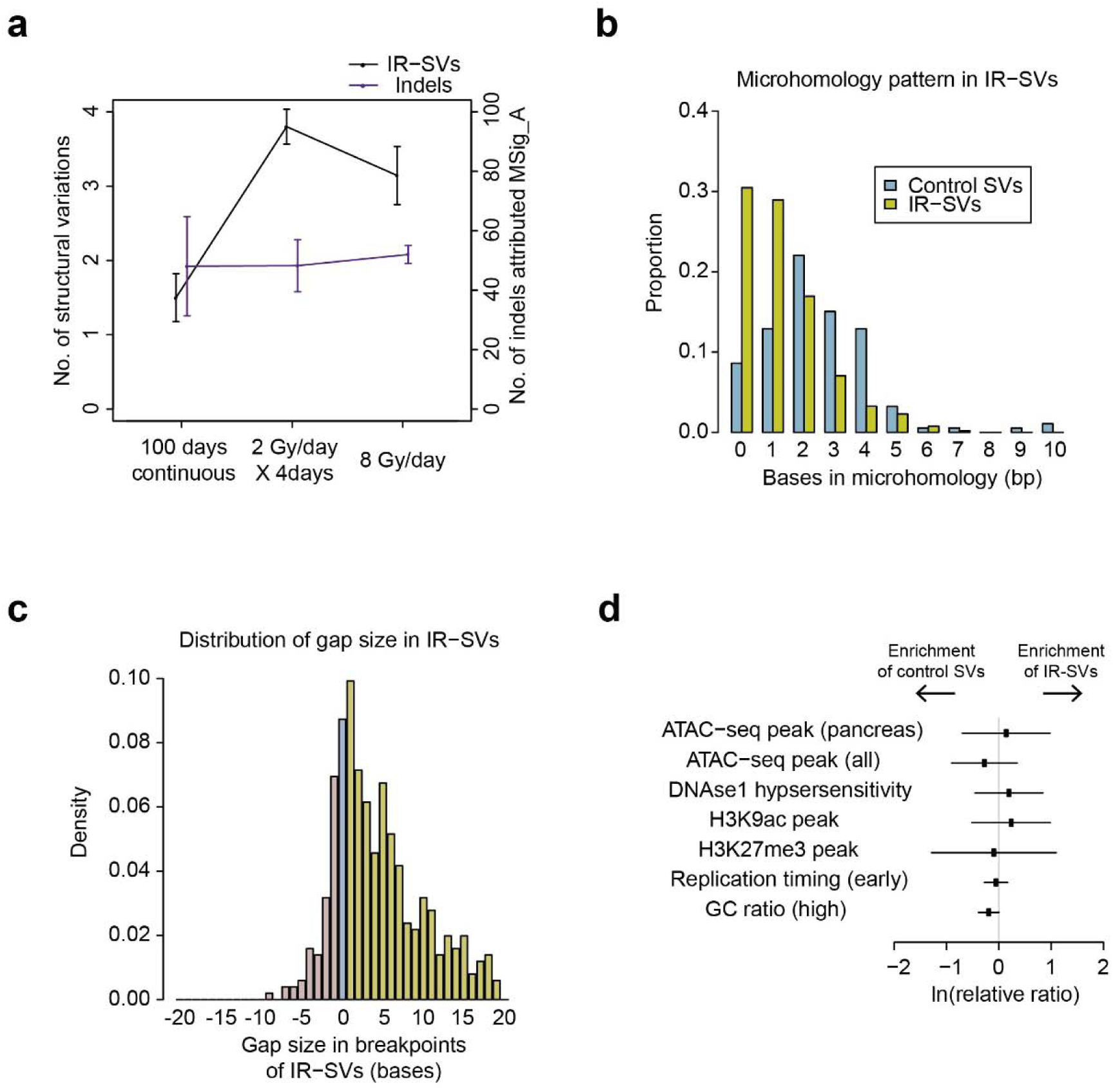
Sequence characteristics of IR-induced structural variations. a. Number of IR-SVs in 8 Gy irradiated organoids with different rates. Organoids irradiated at lower rate (3.3 mGy/hr for 100 days) tend to harbor similar number of indels but lower number of IR-SVs compared with higher rate (0.48 Gy/min). b. Size distribution of microhomology in IR-SVs (yellow). Compared to control SVs (blue), breakpoints of IR-SVs harbor shorter microhomology (*P* < 0.001), which supports that non-homologous end joining is a predominant repair process for IR-induced double strand breaks. c. Size distribution of breakpoint gap in IR-SVs. IR-induced double strand breakages are usually accompanied by a few nucleotide deletions. d. Comparison of control SVs and IR-SVs in relation to genomic contexts. SVs detected in all types of murine organoids were used except ATAC-seq peak (pancreas), in which SVs observed in pancreas organoids were used.

**Extended Data Table 1.**
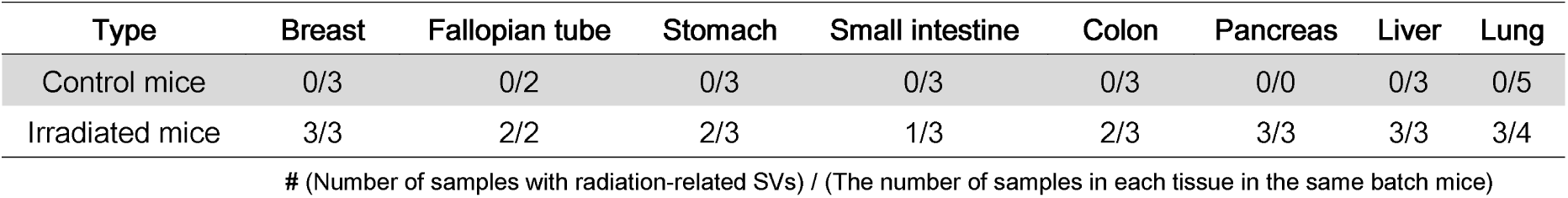
Number of samples harboring IR-SVs in eight tissues. All organoids in this table were derived from 3 control and 3 irradiated (2Gy/day X 4 consecutive days) mice.

