## Supplementary Text for "Mutational impact and signature of ionizing radiation"

**Methods**

***Experimental mice***

Male BALC/c (6-7 weeks old) and female C57BL/6 (8-9 weeks old) mice were purchased from Central Laboratory Animal Incorporated (Seoul, Korea). All experiment protocols were approved by Dongnam Institute of Radiological and Medical Sciences (DIRAMS AEC-2018-007 and DIRAMS AEC-2019-007) and Seoul National University (SNU-180101-1-3). Animal studies were conducted in accordance with guidelines established by the committee on Use and Care of Animals of Dongnam Institute of Radiological and Medical Sciences and Seoul National University Animal Care and Use Committee. Animals were treated humanely in accordance with the Ministry of Food and Drug Safety on the ethical use of animals. Mice were sacrificed with carbon dioxide inhalation euthanasia to get tissues for organoid cultures and spleens for matched germline controls.

***Human normal samples***

For all patients, tumor-adjacent normal tissues or matched blood were collected from irradiated and non-irradiated individuals (controls). The protocol of this study was approved by the institutional review board of Dongnam Institute of Radiological and Medical Sciences (D-1804-023-002 and D-1810-007-002).

Radiation-free human normal colon tissues were acquired from surgical specimen of two colorectal cancer patients (**Supplementary Table 2**). The enrolled patients had no experience of previous chemotherapy or radiotherapy. Tumor adjacent normal colon tissues were cut out from a region >5 cm away from primary tumor mass. Irradiated normal colon samples were taken from a patient who was diagnosed with rectal cancer located 7cm away from anal verge. The patient received preoperative radiotherapy (45 Gy/25 fx + 5.4 Gy/3 fx) with 5-fluorouracil and leucovorin for preserving anus. After ultralow anterior resection surgery, a tumor adjacent normal colon tissue in the radiation field was acquired from a region >5 cm away from primary tumor mass.

Normal human breast samples were collected from adjacent normal tissues from total mastectomy of recurred breast cancer in Dongnam Institute of Radiological and Medical Sciences (**Supplementary Table 2**). Patients had breast conserving surgery 1 (HBIR-1) and 17 (HBIR-2) years ago, followed by adjuvant prophylactic radiotherapy (50 Gy/25 fx + 10 Gy/5 fx as a boost for HBIR-1 and 50.4 Gy/28 fx + 9 Gy/5 fx as a boost for HBIR-2). The HBIR-1 patient received neoadjuvant chemotherapy with Adriamycin, cyclophosphamide and docetaxel at initial diagnosis. The HBIR-2 patient received adjuvant cyclophosphamide, methotrexate and 5-fluorouracil at initial diagnosis. The patient took tamoxifen for several years. After recurrence, the HBIR-2 patient had neoadjuvant chemotherapy with doxorubicin, cyclophosphamide, docetaxel and trastuzumab before total mastectomy.

***Human cancer samples***

After receiving the approval by the institutional review board of Seoul National University Hospital (H-1506-026-678), we collected patients who met the following modified Cahan’s criteria for radiation-induced sarcoma between 2015 and 2019; (A) pathologically confirmed sarcoma which was histologically different from the primary cancer, (B) occurred in the field of radiation, and (3) developed at least 6 months after radiation therapy^1^. Sixteen patients were identified and surgically resected in our institution, and we finally obtained whole genome sequences of ten matched tumor and normal samples. We also collected clinical information including initial tumor diagnosis, radiation dose, latency period, and pathologic diagnosis of radiation-induced sarcoma (**Supplementary Table 2**). Median prescribed radiation dose was 50.4 Gy. Radiation-induced sarcomas were located in extremities (n = 5), trunk (n = 4), and head & neck (n = 1). Radiation-induced sarcomas included 4 undifferentiated pleomorphic sarcoma, 3 undifferentiated spindle cell sarcoma, 2 osteosarcoma, and 1 undifferentiated epithelioid sarcoma.

***Pathology review***

Tumors had been fresh frozen (n = 8) or formalin-fixed/paraffin-embedded (FFPE, n = 2). Normal samples were obtained from blood (n = 7) or adjacent normal tissues of FFPE (n = 1) and fresh frozen specimens (n = 2). Two experienced pathologists reviewed specimens and performed microdissection to isolate tumor and normal tissues.

***Irradiation experiments***

For *in vitro* organoids irradiation, ^137^Cs gamma irradiator (MK1-68; J.L. Shepherd and Associates) was employed. Dose distribution in the floor of irradiation cavity was mapped by 24-well plate containing PBS-submerged radiochromic film (RCF, GAFCHROMIC EBT2, ISP Corporation). Film samples were digitized using a flatbed scanner (Expression 10000XL, Epson) with accompanying software (Silverfast Epson IT8). The optical density from RGB (red, green, and blue) uncompressed tagged film images was analyzed for radiation exposure. After the harvest and dissociation of organoids, the cell suspension was transferred to 1.5 ml Eppendorf tube with culture media. Single cells were irradiated at a dose rate of 31 mGy/s. After the irradiation, the cells were sorted and inoculated in Matrigel (Corning).

For *in vivo* mice irradiation, a cobalt irradiator (GBX200, Best Theratronics) with ^60^Co source (~5,000 Ci) was used. An ionization chamber was used to ensure accurate dosimetry. Ionization chambers for whole-mouse irradiation measured air kerma and an accurate dose was irradiated at an experimental radiation field. The field size of ^60^Co was 20 cm × 15 cm, and the air kerma was measured by inserting the ionization chamber into 50 ml conical tube with holed cap, which was placed at a source-to-chamber distance (SCD) of 60 cm (**Supplementary Fig. 1**). Whole-body γ-ray irradiations of 2 ± 0.098 Gy (0.4763 Gy/min) were carried out in an irradiation room equipped with a ^60^Co source. The standard uncertainty of mice irradiation using cobalt irradiator is shown in **Supplementary Table 9**. Total 8 and 20 Gy were irradiated cumulatively by irradiating 2 Gy per day with 4 times and 10 times, respectively. The animals were sacrificed 24 hours and 2 weeks after irradiation for primary bulk organoid culture and for primary single-cell/crypt culture, respectively. For low-rate long-term irradiation, 8 Gy radiation was irradiated to whole-body with a rate of 3.33 mGy/h using ^137^Cs source (185 GBq) (Chiyoda Technol Corp.)^2^ for 100 days followed by sacrifice after three months from irradiation. Sham mice were also placed in the same tube without irradiation and then sacrificed at the same day as the other irradiated mice.

***Organoid culture***

All organoid establishment procedures and media compositions were adopted from previous literatures with slight modifications. For crypt isolation of murine stomach (antrum)^3^, small intestine^4^ and colon^5^, each tissue was dissected from euthanized mice. Each tissue was opened up longitudinally and washed in cold PBS 3 times. In the case of small intestine, villi were scrapped off using cover slips. Tissues were cut into small 2-4 mm tissues and transferred to 10 mM EDTA (Invitrogen) in 50 ml conical tubes, followed by incubation on an automatic rolling machine for 20-30 min at room temperature. After incubation, the tubes were shaken to extract crypts from the tissues. The supernatant was transferred to new tubes and centrifuged at 300 g for 5 min, and the pellet was washed one time with PBS. Isolated crypts were embedded in Matrigel and plated in a 12- or 24-well plate (TPP). The plates were transferred into an incubator at 37 °C for 5-10 min to solidify Matrigel. Each well was overlayed with 0.5-1 ml of organoid culture media for a 24- or 12-well plate, respectively. Organoid culture media compositions of stomach, small intestine and colon were described in **Supplementary Table 10**.

For mouse pancreas^6^, liver^6^, breast^7^ and fallopian tubes^8^, each tissue was collected from euthanized mice and washed in cold HBSS (Gibco). Each tissue was placed in a 100-mm Petri dish and minced into small pieces using scalpel blades. For pancreas, liver and breast, 10 mg/ml collagenase P (Roche) and 0.1 mg/ml DNAse (Sigma) in HBSS were prepared as digestion solutions. For fallopian tubes, 0.5 mg/ml collagenase I (Sigma) was prepared for digestion. The minced tissue was transferred to 50 ml conical tube and 10 ml of prewarmed digestion buffer was added, followed by incubation at 37 °C with shaking at 230 rpm for 20-45 min. The digested tissues were washed with PBS two times and shaken in PBS. Because insufficiently digested tissues were frequently observed in pancreas and liver, the tubes with digested tissues were allowed to stand for 20-30 seconds to remove the insufficiently digested tissues. Then, supernatant was collected and centrifuged at 300 rcf for 3 min. For breast and fallopian tube, the tubes with digested tissues were centrifuged at 300 rcf for 3 min. After centrifugation, pellets were embedded in Matrigel. Organoid culture media compositions of pancreas, liver, breast and fallopian tubes were described in **Supplementary Table 10**. In the case of liver, isolation media was used for primary culture cases during initial 3-4 days. Then, expansion media was added to culture the liver organoids.

For mouse lung organoid^9^, rib cage was opened up in euthanized mice. Lung perfusion was carried out through right ventricle using 10 ml DPBS in a 10 ml syringe with a 26 G needle. After removing heart, trachea was dissected and cut off at the most proximal portion. Through the proximal opening of the trachea, 2 ml dispase (Corning) was infused until expansion of all lobes of the lung. Whole lung was separated from the thoracic cage with lifting up the trachea. Five lobes were dissected from the whole lung using scissors, followed by digestion in 3 ml PBS with 65 μl of collagenase/dispase (Sigma) in a 37 °C shaking incubator with 230 rpm. After 35 minutes of digestion, 7.5 μl of DNase I (Sigma) was added to the digestion solution. With 10-20 minutes of additional incubation, the digested solution was filtrated using 40 μm strainer (Falcon) to remove insufficiently digested materials. Then, filtrates were centrifuged at 400 rcf for 5 minutes. Because a number of RBCs were frequently observed in the pellet, the pellet was incubated in RBC lysis buffer (Sigma) for 1-1.5 minutes, followed by adding 10 ml of advanced DMEM/F12 and 500 μl of FBS (Gibco) at the bottom. After centrifuging at 400 rcf for 5 minutes, supernatant was discarded, and the pellet was resuspended in 200 μl of PF10 (10% FBS in DPBS). The resuspended solution was incubated with Anti-EpCAM PE antibody (BD Biosciences, 563477), anti-CD31 APC antibody (BD Biosciences, 550274), anti-CD45 APC antibody (BD Biosciences, 561018) and anti-Ly-6A/E APC-Cy7 antibody (BD Biosciences, 560654). After 30-60 minutes incubation, 800 μl PF10 was added and centrifuged at 13,000 rpm for spin-down. The pellet was washed two times with PF10. Then, type 2 alveolar epithelial cells (EpCAM-positive, CD31-negative, CD45-negative and Ly-6A/E-negative) were isolated using fluorescence-activated cell sorter (FACSAria II, BD Biosciences) (**Supplementary Fig. 2**). The type 2 alveolar epithelial cells were embedded in 1:1 mixture of complete lung organoid media (**Supplementary Table 10**)^10^ and Growth Factor Reduced Matrigel (Corning), and the droplets of organoids and Matrigel mixture were seeded in 6.5 mm transwells (Corning) in a 24-well plate. After 20 minutes incubation, 500 ul complete lung media with 10 μM Y-27632 (Sigma) were added into each well.

Plates transferred to humidified 37 °C/5 % CO_2_ incubator and medium was changed every 2-3 days. Organoids were passaged by removing the medium and dissolving Matrigel by adding Cultrex organoid harvesting solution (Trevigen) or cell recovery solution (Corning) for 40-50 minutes on ice. After removing the dissolving solution, 0.5-1 ml prewarmed TrypLE Express (Gibco) or prewarmed Accutase (Stemcell Technology) was added to disrupt organoids, followed by centrifuging at 300 rcf for 3 minutes. In the case of murine breast organoids, 1cc syringe with a 26 G needle was used to dissociate before centrifugation. Pellet was resuspended with Matrigel, and seeded in a new 12- or 24-well plate at ratio of 1:4 to 1:6. After polymerization of Matrigel in a 37 °C incubator, 500-1000 μl complete media were added to each well.

In the several complete organoid media, Wnt3a or R-Spondin-1 conditioned media was included. We prepared the conditioned media using Afamin-Wnt3a producing HEK293 cell line^11^ and Cultrex HA-R-Spondin 1-Fc 293T cell line (Trevigen), respectively. The activity of the harvesting conditioned media was tested using TOP/FOP assay^12,13^.

***Single-cell derived clonal organoid acquisition***

For clonalization of single-cells, we used two alternative strategies. First, bulk organoids were harvested and single-cell dissociated using prewarmed TrypLE Express or Accutase. After centrifuging, supernatants were removed and organoids were resuspended using AdDF+++ (Advanced DMEM/F12 with 10mM HEPES, 1X Glutamax and 1% penicillin/streptomycin). Organoid suspensions were filtered through 40 μm strainer (Falcon). By using a cell sorter (FACSAria II, BD Biosciences), single cells were sorted into a FACS tube with 2 ml of AdDF+++. Single cells were selected in the FACSDiva software on the basis of forward- and side-scatter characteristics^14^. Sorted cells were seeded with Matrigel (500-1500 / well). Alternatively, single whole-crypts were collected from stomach, small intestine, and colon after 14 days from whole-body irradiation. Of note, a crypt is physiologically clonalized *in vivo*, due to the competitive proliferation of single-cells^15^. When mono- to oligo-clonal organoids were acquired right after primary culture, digested materials were sparsely seeded with Matrigel at P0. After growing organoids, a single organoid distant from other organoids was picked up using a 200 μl pipette under microscope. Clonal organoid generation was evaluated by variant allele frequency of somatic point mutations.

***DNA extraction***

Genomic DNA was extracted using DNeasy Blood & Tissue Kit (Qiagen) or TaKaRa MiniBEST Universal Genomic DNA Extraction Kit Ver. 5.0 (TaKaRa Bio Inc.). At the last step, the DNA sample was eluted twice with 40 μl of elution buffer. The concentration of DNA was measured using a Nanodrop 2000 spectrophotometer (Thermo Scientific, ND-2000) and stored at -20 °C or -80 °C before use.

***Whole genome sequencing and read alignment***

DNA libraries for whole genome sequencing (WGS) were generated using Truseq DNA PCR-Free Library Prep Kits (Illumina) from >500 ng of genomic DNA or TruSeq Nano DNA Kit (Illumina) from >100 ng of genomic DNA, according to the manufacturer’s protocols. When genomic DNA was not sufficient to use the PCR-Free Library Kits, Accel-NGS 2S Plus DNA Library Kit for Illumina (Swift) were alternatively used to construct DNA libraries (n = 10; **Supplementary Table 1**). The libraries were sequenced with paired-end (2x151bp) using HiSeq 2500, HiSeq X10, or Novaseq 6000 platforms (Illumina) to generate a minimum of 30X depth for most samples except B3S100 (100X), radiation-induced sarcomas (90X), and tumor-matched normal controls (60X). The raw FASTQ files were aligned to human reference genome, GRCh37/hg19, and mouse reference genome, GRCm38/mm10, using Burrows-Wheeler Aligner mapping tools^16^. Samtools was used to convert sam to bam and to sort^17^. Duplicated reads were marked and removed using the Picard package (<https://broadinstitute.github.io/picard)>. Then, local realignments and base recalibration were proceeded using the Genome Analysis Toolkit^18^.

***Whole transcriptome sequencing of organoids***

Mouse pancreas organoids in Matrigel were irradiated (2 Gy). After irradiation, the irradiated organoids were retrieved from Matrigel at five different time points (0, 0.5 hr, 2 hr, 6 hr, and 24 hr; three biological replicates at each time point). After centrifugation at 300 g for 3 minutes, total RNA was extracted from the organoid pellets using RNeasy Mini Kit (Qiagen, #74104). Total RNA sequencing library was constructed using Truseq Stranded Total RNA Gold kit (Illumina) according to the manufacturer’s protocol.

For validation of *Myc* amplification in murine fallopian tube organoids, total RNA was extracted from FT_8Gy_1 (*Myc* non-amplified; three biological replicates) and FT-8Gy_2 (*Myc* amplified; three biological replicates). TruSeq RNA Library Prep Kit v2 (Illumina) was used to generate cDNA libraries.

We performed high-throughput sequencing with 2 x 101 bp using Hiseq 2500. Adapter contamination was eliminated from the FASTQ files using Cutadapt^19^. Then, the trimmed FASTQ files were aligned to GRCh38/mm10 using STAR v2.6.1d^20^ and normalized counts of RNA expression were calculated using RSEM v1.3.1^21^. The top 2500 genes with highest variations (standard deviations/mean of transcripts per million (TPM) values of each gene) in irradiated organoids at five time points (0, 0.5, 2, 6, and 24 hr) were demonstrated as heatmap (**Fig. 1c**) using R package “ComplexHeatmap”^22^. Enriched pathways were obtained from Enrichr website with input of the top 2500 variable genes^23^.

***Detection of somatic variants***

Strelka2 and Varscan2 were used to find single base substitutions (SBSs) and small insertions and deletions (indels)^24,25^. After taking union of SBS and indel call sets passed in each caller, in-house python script using pysam module was utilized to annotate read information and filter out false positive calls. To exclude sequencing artifacts, loci-specific background mutation rates were calculated using germline samples. In the case of murine samples, normal organoid samples from other mice of the same strains were also applied because the number of germline samples at each strain was not sufficient. This information was further utilized to rule out false positive mutations.

Structural variations (SVs), or genomic rearrangements, were called using Delly 0.7.6^26^. Loci-specific background rearrangement rates were also calculated as described above. Accurate breakpoint positions and microhomology sequences were determined using “SA tag” of clipped reads. Further read information around breakpoints were annotated by in-house script with pysam module, including the number of wild-type read pairs spanning breakpoints with appropriate orientation and normal insert size (≤1000 bp) and the number of variant read pairs spanning breakpoints with inappropriate orientation or large insert size (>1000 bp). Then, breakpoints suggesting false positive variants were removed by in-house script. Briefly, variant calls that meet any of the following criteria were regarded as false positives and excluded from the downstream analysis: (i) breakpoints having high depth of read pairs in a matched germline sample (read pairs ≥ 200), (ii) ≥2 SA tag being the same with a somatic sample at a breakpoint in a matched germline sample, (iii) sequencing artifacts (rearrangements detected in ≥2 unmatched normal samples), or (iv) low supporting variants in somatic samples (less than ten discordant read pairs without supporting SA tag, less than three discordant read pairs with 1 supporting SA tag). Then, all rearrangements were visually inspected using Integrative Genomics Viewer (IGV)^27^ to remove remaining false positive variants and to rescue false negative variants near breakpoints. Gains and losses of <100 bp sequences were classified as indels.

Somatic LINE-1 retrotranspositions were called using TraFiC-mem^28^ and manually investigated using IGV to filter false positive calls. Several false negative calls were rescued during visual inspection of structural variations.

Segmented copy number alterations (CNAs) of clonal organoids were estimated by Sequenza package with modification for GRCm38/mm10 assembly^29^. Except a few organoids with *chromothripsis* or breakage-fusion-bridge cycles, purity and ploidy were assigned as 1 and 2.0, respectively. When abrupt CNAs without breakpoints at callable regions, we manually investigated the regions to find false negative breakpoints using IGV. To show SVs and CNAs in each sample, we used the Circos package^30^.

***Publicly Available Datasets***

In addition to the newly sequenced mouse and human organoids and radiation-induced sarcomas, we obtained additional WGS from previously published cancer datasets to validate mutational signatures of ionizing radiation (IR). For validation of IR-induced indel signatures, 12 radiation-induced cancer^31^, 64 bone sarcoma, and 31 soft tissue sarcoma data^32^ were used. For comparison of short segmental transposition (SST) incidence, 138 lung cancer^33^ and 369 diverse cancer data (liver, esophageal, prostate, bladder, stomach, biliary tract, rectal, and oral cancer, skin cutaneous melanoma) from the PCAWG datasets^32^ were used. Most WGS datasets were re-processed from raw FASTQ files as described above. For 12 radiation-induced tumor samples, we used available variant call sets of SBSs, indels, and SVs for downstream analyses.

***Reconstruction of Genomic Rearrangements.***

Given confident rearrangement call sets (n = 1720) from 156 organoids (**Supplementary Table 11**), we reconstructed the rearrangement patterns and classified them into 16 types of SVs (n = 877), including (i) medium-sized simple-deletion (<1 Mb), (ii) tandem duplication, (iii) templated insertion, (iv) retrotransposition, (v) miscellaneous type, (vi) balanced inversion, (vii) balanced translocation, (viii) long simple-deletion (≥1 Mb), (ix) double minutes, (x) large deletion with local inversion (LDI), (xi) large deletion with translocation (LDT), (xii) short segmental transposition (SST), (xiii) *chromoplexy*, (xiv) *chromothripsis*, (xv) fold-back inversion, and (xvi) others complex rearrangements.

All the balanced inversions and balanced translocations showed no overlapping segments or overlapping less than 10 bp, except one case (225 bp in one 20 Gy irradiated liver organoid). If a pattern of SV suggests balanced inversion or balanced translocation combined with >500 bp copy number loss in one of the breakpoints, we considered those SVs as LDI or LDT. An SV was defined as double minute if one read of a deletion-type rearrangement and one read of a duplication-type rearrangement are located at one breakpoint and the other reads of the two rearrangements are located at another breakpoint together with copy number gain or loss. SST was an ectopic insertion of a short (typically 100-1000 bp) deleted segment like cut-and-paste type mobilization of DNA transposons. *Chromoplexy* was defined as a cluster of reciprocal rearrangements involving three or more chromosomes^34^. *Chromothripsis* is a cluster of localized (usually involving 1 or 2 chromosomes) rearrangements with massive breakpoints exhibiting copy number oscillation between two or three copy number states^35^. Fold-back inversion was defined as a cluster of rearrangements with multiple copy number amplifications, which could be explained by repetitive BFB cycles. Others complex was rearrangements of three or more segments by cut-and-paste mechanism. Some breakpoints (n = 25) were defined as miscellaneous if the rearrangement is not involved in the other 15 types.

In general, complex genomic rearrangements (CGRs) were defined by SVs consisting of ≥3 rearrangements (breakpoints). Simple genomic rearrangements (SGR) were SVs with 1 or 2 breakpoints, including large deletion, balanced inversion, and balanced translocations, and others.

***Monoclonal and polyclonal organoids***

To extract precise mutational burden and signatures harbored in the original single-cell, we excluded polyclonal organoids and subclonal mutations (culture-associated mutations) in an organoid. To filter out polyclonal organoids, the distribution of variant allele frequency (VAF) of somatic SBSs was considered as used in a previous report^36^. Murine organoids with the VAF peak < 0.4 or not FACS-sorted were deemed polyclonal and thus removed from downstream SBS and indel but SV analyses. Of note, all human colon organoid samples were monoclonal and human breast organoids harboring IR-SVs with VAF around 0.5 were regarded as monoclonal organoids.

Of the removed organoids (n = 74), the most (81%) showed a clear VAF peak greater than 0.25, suggesting a dominant clone contributing more than 50% of the cell pool with a few more clones of minor proportions (referred to as ‘dominant-clonal’ organoids). Although the rest 14 organoids (referred to as ‘non-dominant-clonal’ organoids) did not show a meaningfully clear VAF peak, SVs with VAF ≥ ~0.15 were obtained, which indicates that they are not dominated by one clone but still composed of only a handful number of clones. Because we can at least qualitatively examine SVs in these dominant- and non-dominant-clonal organoids, all of the removed organoids were rescued for analyses of IR-SV pattern (**Fig. 2a**).

In monoclonal samples (n = 67 for mouse and n = 15 for human), SBSs and indels with VAF ≥ 0.3 were classified as clonal variants. When we extract SBS and indel signatures (**Figs. 1e-i**), we used clonal mutations in organoid samples. From cancer samples, all somatic SBSs and indels were used to extract mutational signatures (**Fig. 1i**).

***Dose-response curves of ionizing radiation***

To accurately observe dose-response relationship between IR and mutational burden shown in **Figs. 1d**, **4a** and **4b**, we used clonal mutations detected in monoclonal murine pancreas organoids (n = 39), all of which were daughter cells of a single mother cell (PA_0Gy_10).

***Estimation of the numbers of DSBs induced by irradiations***

We estimated the DSBs induced by 1 Gy irradiation using two alternative methods, (1) genome analyses and (2) imaging analyses. From genome analyses, the total number of DSBs is equivalent to the sum of IR-associated breakpoints (including indels and SVs; directly detectable from sequences) and seamless repair counts (directly undetectable). We estimated the seamless repair numbers using breakpoints in SSTs, where end-repairs during junctional ligations were traced. Here, we estimated that about 5% of DSBs were seamlessly repaired and we presumed that the proportion is more or less uniform in other SV types.

Then, the number of DSBs (in the murine genome) was calculated as follows:

$No. of DSBs=1.103/(1-0.05)\times(1.02\times No.of indels+No. DSBs to make SVs)$,

where 1.103 is to adjust for inaccessible genomic regions, 0.05 is the proportion of the seamless repair, and 1.02 is to adjust for repetitive regions where indels are not accurately detected. To extrapolate the estimate to human genome, we used the relative ratio of the sequence length of two reference genomes (1.136; human = 3,101,788,170, mouse = 2,730,871,774 bp).

For imaging analysis, we counted the numbers of γ-H2AX foci in irradiated organoids combined by super-resolution imaging technique. The details are independently described below.

***Mutational Signature analysis of single base substitutions and indels***

We borrowed the conventional presentation of mutational signature from the COSMIC database^37^, using a feature set that summarizes SBS and indel variations by 96 and 83 types of mutational events, respectively^38^.

To learn new mutational signatures from a set of samples, we referred to the original MATLAB code of the current SigProfiler, one of the widely used tools for signature extraction, published in 2013^39,40^. Overall, we followed the same procedure, except for manually choosing some of the computation and model parameters. In particular, we took great care in determining $k$, the number of presumed mutational processes, mainly through examining whether a set of signatures obtained from the chosen $k$ leads to a coherent story without under- or over-explaining the mutational history. We also used a measure of stability and reconstruction error to get a reasonable initial guess for $k$^39^.

In the given methodology, learning new mutational signatures amounts to carrying out a mathematical technique called non-negative matrix factorization (NMF). Briefly speaking, we are given with an input matrix $V\in R^{m\times n}$ (i.e., $m$ features and $n$ samples) and our objective is to solve $V=WH+\varepsilon$ for $W$ and $H$ where $W\in R_{+}^{m\times k}$, $H\in R_{+}^{k\times n}$, and an error term $\varepsilon\in R_{+}^{m\times n}$. We seek for an approximate solution by formulating it as an optimization problem:

$\underset{W, H}{\mathrm{argmin}} D(V||WH)$ where $D(V|\left| WH \right)=\sum_{\mathrm{ij}} \left( V_{\mathrm{ij}}\log\frac{V_{\mathrm{ij}}}{\left( WH \right)_{\mathrm{ij}}}-V_{\mathrm{ij}}+\left( WH \right)_{\mathrm{ij}} \right)$,

where $D(V||WH)$ is a generalized Kullback-Leibler divergence between $V$ and $WH$^41^. To avoid overfitting and promote stability of a solution, a bootstrap aggregating is used, where K-means clustering is used to obtain an averaged solution^39^.

We constructed 4 data sets: human SBS, mouse SBS, human indel, and mouse indel data. The human SBS and indel data came from 15 normal organoids and 22 radiation-induced cancers. The mouse SBS and indel data came from 67 monoclonal organoids. We extended the mouse data by replicating 67 samples that we can infer clonal mutations. For the added samples, we filtered out variants where VAF < 0.3, in order to get clonal mutations and exclude mutations irrelevant to radiation exposure.

For further validation, we constructed 4 extended data sets that include the data above. The extended human SBS data includes 2 replicates of 15 organoids (clonal samples: VAF ≥ 0.3; raw samples: 1 ≥ VAF ≥ 0), 22 radiation-induced cancers, and 95 PCAWG sarcoma samples. The extended human indel data includes the same data as the SBS data except that 24 out of 95 PCAWG samples are excluded because of low indel calls (< 60 indels, considered noisy). The extended mouse SBS data includes 2 replicates of 67 monoclonal organoids (clonal samples: VAF ≥ 0.3; raw samples: 1 ≥ VAF ≥ 0) and the other 74 polyclonal organoids.

Of a note, we noticed that some samples had peculiar mutational spectra that are distinct from the majority of the others. Inclusion of these samples led to a greater number of mutational signatures. However, the additional signatures appeared to be due to a batch effect, found only in a single or a group of samples that share common attributes (e.g., coming from the same facility where the same library preparation was done, etc.). Nevertheless, we did not exclude these samples for completeness. As a side effect, these singular signatures happened to distribute its mass in a mutational spectrum to irrelevant samples. In other words, in the resulting mutational spectrum, they appear to take part in the samples in which they are actually not likely to be present. This is an undesirable but known behavior of the NMF algorithm. To make the analysis simple, we ignored it because it only mildly does so and does not affect our conclusion.

After the analysis of learning new mutational signatures, we wanted to assess to what extent the prior knowledge can explain the observed mutational spectra. We fitted the known COSMIC signatures to individual samples in the same data set. Fitting known signatures to a sample requires a slightly different consideration but can be done essentially in the same way as before (i.e. the same optimization problem). The only difference is that we now fix $W$ to be a constant matrix instead of letting it be variable and do not perform bootstrap aggregating. To construct $W$, we used version 3 of the COSMIC mutational signatures.

To accommodate inflated proportions of unexplained mutations from using the fixed prior $W$, we defined “unexplained” counts to quantify to what extent the prior cannot explain the observed mutations. For that, we regarded each mutational signature as a multinomial distribution and mutations as samples from a convex combination of the distributions. The residual is then regarded as a deviation from the mean that arises from random sampling. The rationale is that residuals that do not fall within a certain expected interval should be attributed to unknown mutation-generating processes. More precisely, given that fitting on a sample resulted in the residual $r=v-\hat{v}$ where $\hat{v}$ is our reconstructed mutational spectrum, we define an unexplained count for the sample as

$$\sum_{i}^{m} \max\left( 0,1_{r_{i}\geq0}\left( \left| r_{i} \right|-\left| \text{CI}_{95\%}^{+}\left( \hat{v_{i}} \right) \right| \right)+1_{r_{i}<0}\left( \left| r_{i} \right|-\left| \text{CI}_{95\%}^{-}\left( \hat{v_{i}} \right) \right| \right) \right),$$

where $m$ is the number of feature (i.e., 96 for SBS and 83 for indel) and $1_{C}(x)$ is an indicator function that becomes $x$ only when the condition $C$ is met and 0 otherwise. $\text{CI}_{95\%}^{+}\left( \hat{v_{i}} \right)$ (or $\text{CI}_{95\%}^{-}\left( \hat{v_{i}} \right)$) is a value obtained by subtracting $\hat{v_{i}}$ from the upper (or lower) bound of 95% confidence interval of $\hat{v_{i}}$. We assumed that $\hat{v_{i}}$ is a reasonable mean estimate of the sum of binomial random variables and used Jeffreys method to compute the binomial confidence interval^42,43^. Our definition may underestimate unexplained counts that may potentially be attributable to unknown mutational processes. It can be accounted for by the binomial sum variance inequality ^44^ and the fact that we ignore counts that fall within the 95% CI of $\hat{v_{i}}$. Nevertheless, our definition is conservative in the sense that when the unexplained count starts to increase we can be more sure than the other way around (i.e., weighted counting within the interval) about the presence of potentially unknown processes.

***Assay for Transposase-Accessible Chromatin using sequencing (ATAC-seq) analysis***

To investigate whether chromatin status was associated with the formation of IR-SVs, we performed ATAC-seq on control murine pancreas organoids (n = 4, biological replicates). According to the published protocol^45^, nuclei isolation was initially performed as follows. Organoids (approximately 10k-100k cells) were retrieved from Matrigel using Cell Recovery Solution (Corning). After centrifugation at 300 g for 3 minutes, 10k-100k cells were prepared, and resuspended in 100 μl cold lysis buffer (Tris·HCl pH7.4 10 mM, NaCl 10 mM, MgCl2 3 mM, BSA 1%, Tween-20 0.1%, NP40 0.1% (Sigma), Digitonin 0.01% in distilled water). After 3 minutes incubation, 1 ml wash buffer (Tris·HCl pH7.4 10 mM, NaCl 10 mM, MgCl2 3 mM, BSA 1%, Tween-20 0.1% in distilled water) was added and centrifuged at 500 g for 5 minutes. Supernatant was discarded and nuclei was resuspended in 95 μl DPBS. Approximately 10-20k nuclei were prepared and centrifuged at 500 g for 5 minutes. The supernatants were discarded and remaining nuclei were gently resuspended in 16.5 μl of DPBS.

After the nuclei isolation, transposition reaction mixture (16.5 μl nuclei in DPBS, 12.5 μl 2x TD buffer (Illumina), 0.5 μl 1% digitonin, 0.5 μl 10% tween-20, and 2.5 μl transposase (Illumina) in 5 μl distilled water) was prepared and incubated at 37 °C for 45 minutes. Post clean-up process was performed using MinElute PCR Purification Kit (Qiagen), and the transposed DNA was eluted in 10ul Elution Buffer. PCR amplification of the transposed DNA was performed with the following in PCR tubes (10 μl of transposed DNA, 2.5 μl Nextera index primer1 (Illumina), 2.5 μl Nextera index primer2, 25 μl NEBNext High-Fidelity 2x PCR Master Mix (New England Labs) in 10 μl distilled water). Thermal cycle was as follows: 72 °C 5 minutes, 98 °C 30 seconds, 5 cycles of (98 °C 10 seconds, 63 °C 30 seconds, 72 °C 60 seconds), then 4 °C hold.

To determine the appropriate number of qPCR cycles, 5 μl of PCR amplified DNA was run with 5 μl NEBNext High-Fidelity 2x PCR Master Mix, 0.5 μl Nextera index primer1, 0.5 μl Nextera index primer2 in 3.85 μl distilled water + 0.15 μl SYBR Green I (Invitrogen). A cycle was as follows: 98 °C 30 seconds, 20 cycles of (98 °C 10 seconds, 63 °C 30 seconds, 72 °C 1 minutes). The number of cycles corresponds to 1/3 of maximum fluorescent intensity was applied to qPCR for a 45 μl remnant of the PCR amplified DNA. Lastly, we performed double library size selection using SPRIselect. Briefly, 22.5 μl (0.5x) of SPRIselect beads were added, and supernatant was transferred. Then, 58.5 μl (1.8x) of SPRIselect beads were added, and libraries attached to the beads were eluted in 40 μl of Elution Buffer. The final product (2x101 bp) was sequenced by Hiseq 4000 (Illumina).

Using the raw fastq files of control organoids (n = 4), we performed primary data processing according to the ENCODE ATAC-seq Pipeline (<https://github.com/ENCODE-DCC/atac-seq-pipeline>) including alignment to GRCm38/mm10 by Bowtie2 aligner ^46^. Then, peak calling was done by Model-based Analysis for CHIP-Seq (MACSv2, <https://github.com/macs3-project/MACS>). We used narrow peak calls in the downstream analysis.

***Association of IR-SV incidences with genomic features***

To investigate whether IR-induced DSBs occurred more frequently in a certain genomic context, we first collected several genomic contexts information from our own experiment (i.e. ATAC-seq in murine pancreas organoids) as well as from Mouse ENCODE, including DNase-seq of liver, H3K9ac CHIP-seq of liver, H3K27me3 CHIP-seq of liver (8 weeks old male C57BL/6), and Repli-chip of embryonic fibroblast (13.5 days male C57BL/6 embryo)^47^. GC ratio in 50 bp windows was calculated using mouse reference genome sequence. If genomic coordinates were from mm9, we converted them to mm10 using liftOver^48^. Using SVs identified from the irradiated murine organoids (IR-SVs) and SVs from controls (SVs from non-irradiated murine organoids), we tested enrichments of IR-SVs in euchromatin (open chromatin), early replicating regions, or genomic regions with high GC ratio, using the proportions of breakpoints around regions spanning peaked position ± 20 bp. For ATAC-seq data, we additionally tested enrichment of IR-SVs detected in murine pancreas organoids.

***Sanger*** ***validation of short segmental transpositions (SSTs)***

For SST validation, primers were designed to cover the candidate deletion and insertion area of the SSTs:

PA_2Gy_11 deletionF, 5’-GCCTGTGTTCAAATTGGGGG-3’;

PA_2Gy_11 deletionR, 5’-AAAACCCATGCCCTCTGCTT-3’;

PA_2Gy_11 insertionF, 5’-CCACATGGAACCTTATGCTGC-3’;

PA_2Gy_11 insertionR, 5’-TTCTGACTGCCTTGGCACAG-3’;

These regions were PCR-amplified using 500 ng genomic DNA, 5× buffer 10 μl, 0.25 mM of each dNTP, 0.2 μM primers and 0.5 unit of PrimeSTAR HS Taq polymerase (Takara) in a final volume of 50 μl or 500 ng genomic DNA, 10× buffer 5 μl, 0.2 mM of each dNTP, 0.2 μM primers and 1.25 unit of Pfu DNA polymerase (ThermoFisher) in a final volume of 50 μl at 95 °C for 10 min; followed by 30 cycles of 98 °C for 10 s, 60 °C for 10 s, 72 °C for 30 s; with a final extension at 72 °C for 10 min (Taq polymerase) or 98 °C for 5 min; followed by 98 °C for 10 s, 60 °C for 30 s, 72 °C for 30 s; with a final extension at 72 °C for 7 min (Pfu polymerase). The PCR reaction was run in the Veriti Thermal Cycler (Applied Biosystems) according to the manufacturer’s protocol. The PCR products were cloned into pTOP Blunt V2 vector system (Enzynomics) and Sanger sequencing was carried out using M13 universal primer.

***Viability*** ***assay***

Dissociated pancreas organoids were irradiated (0, 1, 2, 4, and 8 Gy; three biological replicates at each dose) and re-seeded in a 24-well plate (25,000/well). Growth of irradiated organoid was visually monitored up to 6 days. At day 6 after plating, organoids were harvested for quantitative viability assay using CellTiter-Glo 3D kit (Promega). Total ATP amount in each well was measured by luminometer (1420 Victor Light).

***γ-H2AX immunostaining and 3D visualization***

Mouse pancreas organoids on 12 well plates were irradiated (0 Gy and 2 Gy) and incubated for 1 hour at 37 °C. Then the organoids were harvested into 15 ml tube without dissociation procedure and centrifuged at 500 g for 3 min. Collected organoids were fixed in 4% formaldehyde for 20 min and permeabilized with 0.5% Triton X100 in PBS for 20 min at room temperature. Blocking was performed with 4% bovine-serum albumin (BSA) in buffer (0.2% Triton X-100, 0.05% Tween 20 in PBS) for 1 hour. The blocked samples were incubated with a primary antibody (phospho-histone H2A.X (Ser139) antibody, Cell signaling, #2577) in PBST (PBS with 0.1% (w/v) Triton X-100 and 0.02% (w/v) sodium azide) with a 1:300 dilution at 37 ºC for 4 h, followed by washing at 37 ºC for 2 hours in PBST three times. The samples were then incubated with a secondary antibody (donkey anti-rabbit IgG antibody-Alexa Fluor Plus 488, Invitrogen, #A32790) in PBST with a 1:300 dilution at 37 ºC for 2 hours, followed by washing at 37 ºC for 1 hour in PBST three times. For nuclear staining, the samples were incubated in 1 ug/ml DAPI (Invitrogen) in PBST for 30 min followed by brief PBST washing three times. The samples were mounted on a slide glass with a spacer (iSpacer 0.15 mm, SUNJin Lab) filled with PBS and then covered by a coverslip. The mounted samples were imaged by confocal laser-scanning microscopy (Zeiss LSM 780) with a 63x objective (C-Apochromat 63x/1.20 W Korr M27).

***Physical expansion of organoids for super-resolution imaging***

For super-resolution microscopy of organoids, we used a modified magnified analysis of proteome (MAP) protocol^49^. Organoids were embedded in dense polyelectrolyte hydrogels via a physical tissue-gel hybridization approach^50^, for which a modified MAP solution with a decreased concentration of acrylamide was used (20 % (w/v) acrylamide (Sigma), 10 % (w/v) sodium acrylate (AK Scientific), 0.1 % (w/v) bis-acrylamide (BIORAD), 0.03 % (w/v) VA-044 (Wako Chemical) in PBS). Immunolabeled organoid samples were post-fixed with 4% PFA in PBS for 10 min followed by brief PBST washing three times. Post-fixed organoid samples were incubated in the modified MAP solution for 2 hours. In order to locate organoids efficiently in 100-um-thick gels in which organoids were embedded, we used a 3 mm punch to take out regions of interest containing organoids. After expanding the 3 mm punched gel with distilled water three times, about a 3.2ⅹ linear expansion ratio was observed. Samples were mounted on a slide glass with distilled water by using the Blu-Tack adhesive (Bostik) as a spacer, sealing with a coverslip on the top. The mounted samples were imaged using the confocal laser scanning microscope with the 63x objective.

**γ-H2AX counting**

ImageJ^51^ was used for image handling. We divided the microscopic fields of view into 3-by-3 to select cells in organoid images. Fifty cells, recognized through the nuclear staining using DAPI, were randomly selected to count the numbers of γ-H2AX foci found in the Alexa Fluor Plus 488 channel. When selecting the cells, those with broad and high intensity signals of γ-H2AX were avoided to minimize error. The number of DSBs in each selected cell were manually counted.

***Statistics***

All statistical calculations were performed with R version 3.6.0 (R Core Team, Vienna, Austria)^52^. Linear regression (lm function with “qr” method) was used to show significant relationship between two continuous variables. For comparison of mutational signature exposures in control and irradiated samples, Quasi-Poisson (glm function with “quasipoisson” family) test was applied.

For the other analysis, we generally used Pearson’s chi-square test for categorical variables, Wilcoxon rank sum test for non-parametric continuous variables, and t-test for parametric continuous variables. *P* < 0.05 (two-tailed) was considered statistically significant.

**Supplementary Fig. 1. Schematic illustration of *in vivo* whole-body irradiation**

Mice were placed in a 50 ml tube and then 60 cm away from the radiation source. SCD, Source-to-Chamber Distance.

**Supplementary Fig. 2. Representative FACS plot for isolation of murine type II pneumocytes.**

**Supplementary Tables**

**Supplementary Table 1.** Characteristics of 156 control and irradiated normal organoids derived from mice and humans.

**Supplementary Table 2.** Clinical characteristics of human samples.

**Supplementary Table 3.** Numerical matrices of mouse SBS signatures according to the 96 contexts.

**Supplementary Table 4.** Numerical matrices of human SBS signatures extracted from human organoids and radiation-induced tumors.

**Supplementary Table 5.** Numerical matrices of mouse indel signatures according to the 83 contexts.

**Supplementary Table 6.** Numerical matrices of human indel signatures extracted from human organoids and radiation-induced tumors.

**Supplementary Table 7.** Numerical matrices of the supplementary human indel signatures extracted from human organoids, radiation-induced tumors, and 71 PCAWG sarcoma samples.

**Supplementary Table 8.** Number of indels attributed to each human supplementary indel signature in 15 human organoids, 22 radiation-induced secondary malignancies, and 71 PCAWG sarcoma samples.

**Supplementary Table 9**. Mice irradiation uncertainty

**Supplementary Table 10**. Media compositions of organoid cultures used in this study.

**Supplementary Table 11**. List of rearrangements in 58 control and 98 irradiated organoids. Period: no specified loci, usually repetitive regions.

37	Catalogue Of Somatic Mutations In Cancer (COSMIC), <https://cancer.sanger.ac.uk/cosmic/signatures>

38	Catalogue Of Somatic Mutations In Cancer (COSMIC), <https://cancer.sanger.ac.uk/cosmic/signatures>

39	Alexandrov, L. B., Nik-Zainal, S., Wedge, D. C., Campbell, P. J. & Stratton, M. R. Deciphering signatures of mutational processes operative in human cancer. *Cell reports* **3**, 246-259, doi:10.1016/j.celrep.2012.12.008 (2013).

40	Alexandrov, L. B. *et al.* The repertoire of mutational signatures in human cancer. *Nature* **578**, 94-101, doi:10.1038/s41586-020-1943-3 (2020).

41	Lee, D. D. & Seung, H. S. Learning the parts of objects by non-negative matrix factorization. *Nature* **401**, 788-791 (1999).

42	Brown, L. D., Cai, T. T. & DasGupta, A. Interval Estimation for a Binomial Proportion. *Statistical Science* **16**, 101-117 (2001).

43	Young, D. S. tolerance: An R Package for Estimating Tolerance Intervals. *Journal of Statistical Software; Vol 1, Issue 5 (2010)* (2010).

44	Hoeffding, W. On the Distribution of the Number of Successes in Independent Trials. *Ann. Math. Statist.* **27**, 713-721, doi:10.1214/aoms/1177728178 (1956).

45	Buenrostro, J. D., Wu, B., Chang, H. Y. & Greenleaf, W. J. ATAC-seq: A Method for Assaying Chromatin Accessibility Genome-Wide. *Current protocols in molecular biology* **109**, 21.29.21-29, doi:10.1002/0471142727.mb2129s109 (2015).

46	Langmead, B. & Salzberg, S. L. Fast gapped-read alignment with Bowtie 2. *Nature Methods* **9**, 357-359, doi:10.1038/nmeth.1923 (2012).

47	Davis, C. A. *et al.* The Encyclopedia of DNA elements (ENCODE): data portal update. *Nucleic Acids Res* **46**, D794-d801, doi:10.1093/nar/gkx1081 (2018).

48	Kent, W. J. *et al.* The Human Genome Browser at UCSC. *Genome Res* **12**, doi:10.1101/gr.229102. Article published online before print in May 2002 (2002).

49	Ku, T. *et al.* Multiplexed and scalable super-resolution imaging of three-dimensional protein localization in size-adjustable tissues. *Nature Biotechnology* **34**, 973-981, doi:10.1038/nbt.3641 (2016).

50	Ku, T. *et al.* Elasticizing tissues for reversible shape transformation and accelerated molecular labeling. *Nature Methods* **17**, 609-613, doi:10.1038/s41592-020-0823-y (2020).

51	Abramoff, M., Magalhães, P. & Ram, S. J. Image Processing with ImageJ. *Biophotonics International* **11**, 36-42 (2003).

52	R Core Team (2019). R: A language and environment for statistical computing. R Foundation for Statistical Computing, Vienna, Austria. URL <https://www.R-project.org/>.
