## Supplementary figures and images for "Mutational impact and signature of ionizing radiation"

### Supplementary_Figure_1

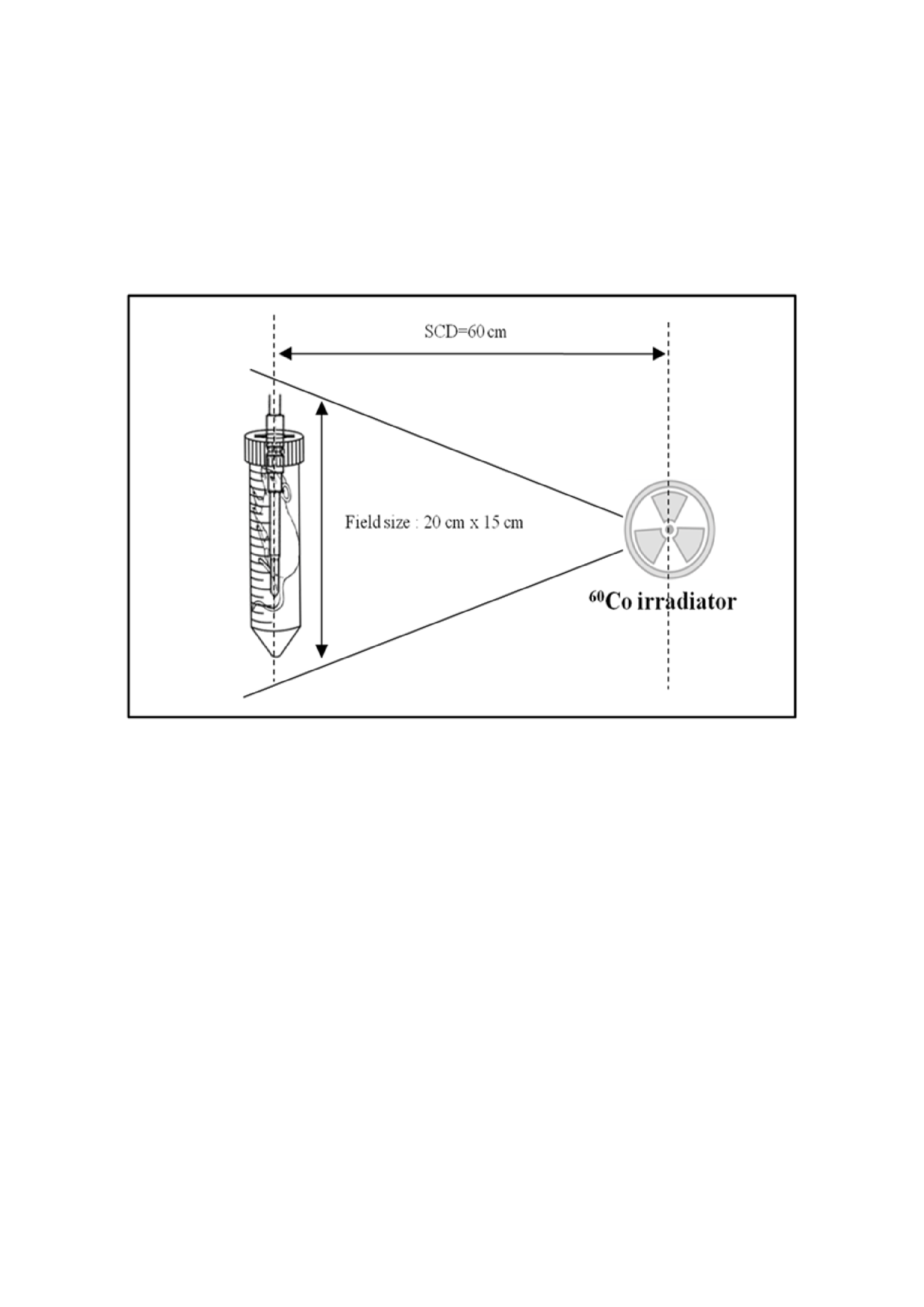

### Supplementary_Figure_2

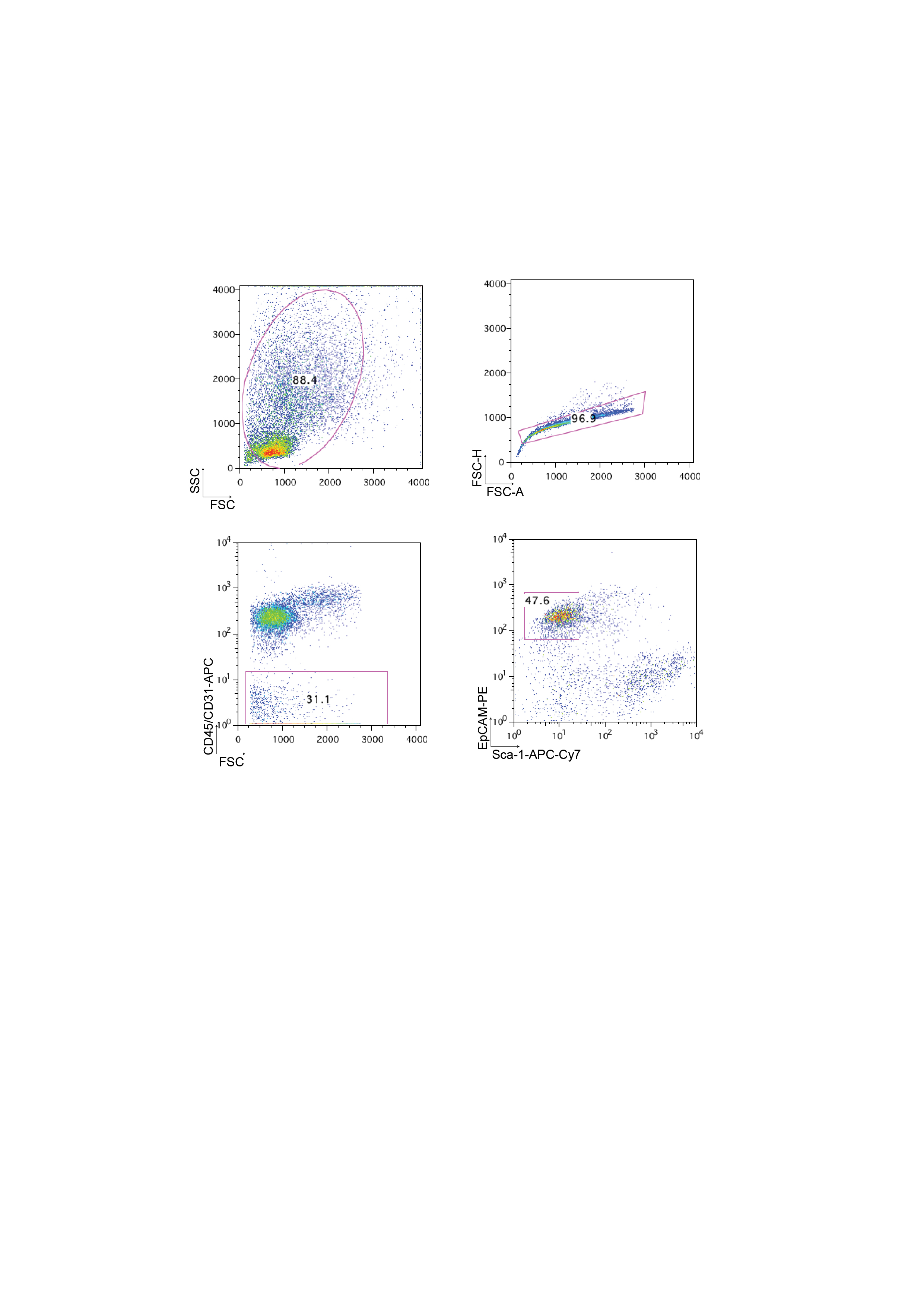
